# RNA polymerase II dynamics and mRNA stability feedback scale mRNA in proportion to cell size

**DOI:** 10.1101/2021.09.20.461005

**Authors:** Matthew P. Swaffer, Georgi K. Marinov, Huan Zheng, Crystal Yee Tsui, Andrew W. Jones, Jessica Greenwood, Anshul Kundaje, William J. Greenleaf, Rodrigo Reyes-Lamothe, Jan M. Skotheim

## Abstract

A fundamental feature of cellular growth is that protein and RNA amounts scale with cell size so that concentrations remain constant. A key component to this is that global transcription rates increase in larger cells, but the underlying mechanism has remained unknown. Here, we identify RNAPII as the major limiting factor increasing transcription with cell size in budding yeast as transcription is highly sensitive to the dosage of RNAPII but not to other components of the general transcriptional machinery. Our experiments support a dynamic equilibrium model where global transcription at a given size is set by the mass-action recruitment kinetics of unengaged nucleoplasmic RNAPII, and DNA content. This drives a sub-linear increase in transcription with size, which is precisely compensated for by a decrease in mRNA decay rates as cells enlarge. Thus, limiting RNAPII and feedback on mRNA stability work in concert to ensure mRNA concentration homeostasis in growing cells.

## INTRODUCTION

Cell size has a profound impact on cellular physiology because it sets the scale of subcellular structures, metabolism, surface-to-volume ratios, and, most crucially, cellular biosynthesis. Biosynthesis scales with cell size so that total RNA and protein amounts increase in proportion to cell size and concentrations remain approximately constant as a cell grows (**Fig. 1A**). Nuclear concentrations are also expected to be constant because the nuclear volume increases in proportion to cell volume (Jorgensen et al., 2007; Neumann and Nurse, 2007). Thus, by scaling RNA and protein amounts with their size, cells maintain approximately constant concentrations of key enzymes and reactants supporting core cellular processes. A key aspect of this biosynthetic scaling is that global transcription rates increase in larger cells (Elliott, 1983a, b; Elliott and McLaughlin, 1979; Fraser and Nurse, 1978, 1979; Padovan-Merhar et al., 2015; Sun et al., 2020; Zhurinsky et al., 2010) (**Fig. 1B**). This size-dependent transcriptional scaling is thought to ensure constant concentrations of global mRNA, rRNA and tRNA, which drives increased protein synthesis in proportion to cell size.

**Figure 1.**
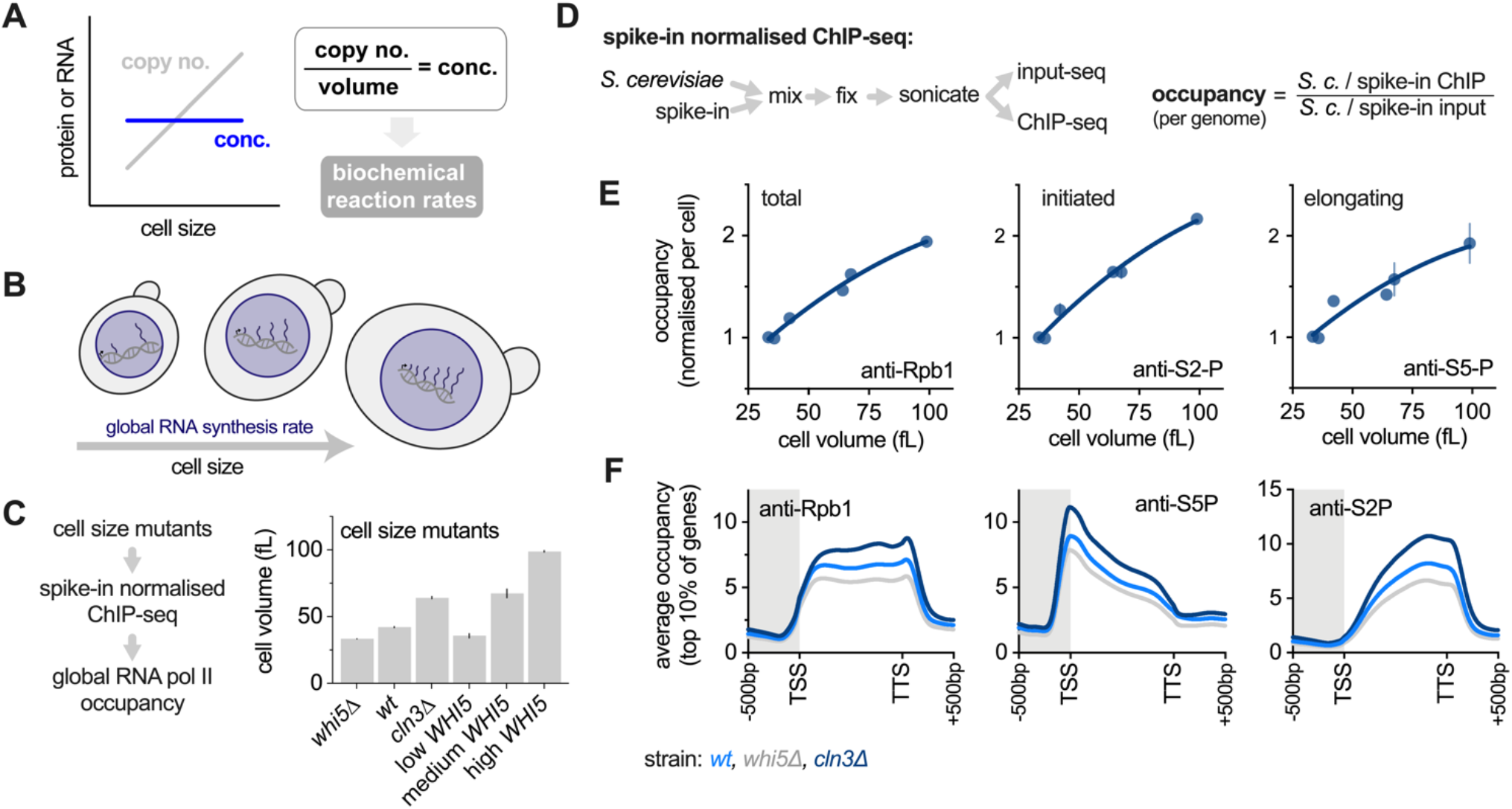
More RNAPII is loaded onto the genome with increasing cell size. See also Fig. S1-S2 **(A)** Macromolecule copy number increases in proportion to cell size to keep RNA and protein concentrations constant. **(B)** Global RNA synthesis rates increase with cell size. **(C)** Mean cell volume determined by Coulter counter of cell size mutants used to in (E-F). Low, medium, and high *WHI5* expression is from a dose-responsive beta-estradiol induced promoter (see also Fig. S2A-B). **(D)** Schematic illustrating the workflow for spike-in normalized ChIP-seq and how the global relative occupancy is calculated (see methods and Fig. S1 for details). **(E)** The occupancy per cell of total Rpb1, initiated Rpb1 (anti-S5-P) and elongating Rpb1 (anti-S2-P) in the size mutants shown in (D) plotted as a function of cell size. Each point shows the mean (±range) of two biological replicates. **(F)** Average occupancy across the gene bodies of the top 10% of genes for total Rpb1, initiated Rpb1 (anti-S5-P), and elongating Rpb1 (anti-S2-P) in *WT, whi5Δ, and cln3Δ* cells. See Fig. S2C for low, medium and high *WHI5* conditions. Mean of two biological replicates is shown.

The importance of this biosynthetic scaling is highlighted by the observation that scaling only occurs within the physiological range of cell sizes. Beyond this range, both RNA and protein amounts cannot keep pace with the expanding cell volume, their concentrations start to decrease, and the cytoplasm becomes progressively diluted (Neurohr et al., 2019; Zhurinsky et al., 2010). This break down in biosynthesis is associated with a dramatic decline in many aspects of cellular physiology and increasing evidence points to this being a causal driver of cellular ageing and senescence (Cheng et al., 2021; Lanz et al., 2021; Lengefeld et al., 2020; Neurohr et al., 2019).

While it is now clear that biosynthetic scaling with cell size is essential for maintaining optimal cellular functionality, the mechanistic origin of RNA scaling has remained mysterious, despite being first reported by foundational radio-labelling experiments in the 1970s (Elliott, 1983a, b; Elliott and McLaughlin, 1979; Fraser and Nurse, 1978, 1979). This is because transcription increases continuously as a cell grows, even among cells with the same amount of template DNA. To explain this phenomenon several theoretical models have been proposed based on a hypothesised ‘limiting factor’ (Lin and Amir, 2018; Marguerat and Bahler, 2012; Padovan-Merhar et al., 2015; Sun et al., 2020; Zhurinsky et al., 2010). In these models, there is a limiting factor that increases in amount in proportion to cell size so that its concentration is constant. Crucially, it is the amount of this limiting factor, rather than its concentration, that is proposed to set the global transcription rate because it is titrated against the genome so that, for example, similarly sized diploid and haploid cells will transcribe at a similar total rate (Lin and Amir, 2018; Sun et al., 2020). While such a limiting factor model could in principle underlie the size-scaling of transcription, there has been no direct experimental test of this model and no empirical identification of the requisite limiting factor.

Here, we show that more transcriptional machinery is loaded onto the genome as cell increase in size and we identify RNA polymerase II (RNAPII) as the major limiting factor for increasing transcription with cell size in budding yeast. In contrast, other components of the transcriptional machinery are not limiting and the chromatin environment into which transcriptional machinery is loaded is effectively constant regardless of cell size. Importantly, our data disprove the previously proposed titration models and instead support a dynamic equilibrium model that accurately predicts RNAPII loading at a given size as a function of the mass action binding kinetics between the free nucleoplasmic RNAPII pool and the genome. However, this only results in a sub-linear increase of transcription that is not directly proportional to cell size. This means that to maintain approximately constant global mRNA concentrations an additional feedback mechanism on mRNA decay rates is required, which we subsequently validate experimentally. Thus, the scaling of mRNA amounts with cell size is driven by both the mass action recruitment kinetics of RNAPII to the genome and feedback regulation on mRNA stability.

## RESULTS

### RNA polymerase II (RNAPII) occupancy increases with cell size in budding yeast

Previous observations that bulk transcription rates increase with budding yeast cell size (Elliott and McLaughlin, 1979) suggest that larger cells should have more RNAPII loaded on their genomes than smaller cells. To test this, we employed a spike-in normalized ChIP-seq methodology to measure RNAPII occupancy (**Fig. 1C-D**). While conventional ChIP-seq only resolves differences in the relative binding at different genomic loci, spike-in normalization also measures systematic differences in the amount of DNA bound protein (*i.e.*, the relative occupancy) (Hu et al., 2015). First, we performed controls to confirm that this approach is quantitative across a wide dynamic range and is robust to variation in the spike-in mixing ratio and cell density (**Fig. S1**).

We then measured global RNAPII occupancy in budding yeast cultures of different average cell sizes, but with similar growth rates (**Fig. 1C & S2A-B**). This shows that the total amount of RNAPII loaded on the genome increases with cell size (**Fig. 1E**). The initiated and elongating RNAPII populations, identified by S5 and S2 phosphorylation in the C-terminal heptapeptide repeats on Rpb1, also increased similarly in larger cells (**Fig. 1E**). Moreover, the relative distributions across gene bodies of total, initiated (S5-P), and elongating (S2-P) RNAPII increase uniformly with cell size (**Fig. 1F**). Taken together, these data indicate that the initiation rate of RNAPII is likely to be primarily responsible for its size-dependent increase on the genome (Sun et al., 2020).

### Total and chromatin bound transcriptional machinery amounts increase with cell size

The increase in transcription with cell size has previously been attributed to a hypothetical limiting component of the transcriptional machinery (Lin and Amir, 2018; Padovan-Merhar et al., 2015; Sun et al., 2020; Zhurinsky et al., 2010). In such models, the amount of the limiting factor increases in proportion to cell size resulting in its increased binding to the genome to drive more polymerase loading and increased transcription in larger cells. In principle, any factor essential for transcription could be such a limiting factor. While this hypothesis has been repeatedly discussed in the literature, it has not been experimentally tested and, crucially, there has been no empirical identification of the requisite limiting factor(s).

To identify candidates for the limiting factor, we first analysed the protein scaling of RNAPII and each of the general transcription factor complexes, which together constitute the RNAPII pre-initiation complex (PIC). The protein amounts of all PIC components that we examined increased in close proportion to cell size consistent with the predicted property of a limiting factor (**Fig. 2A**). We next sought to determine how these factors change their association with chromatin in larger cells. To this end, we adapted a chromatin purification technique called ChEP (<u>Ch</u>romatin <u>E</u>nrichment for <u>P</u>roteomics) (Kustatscher et al., 2014) to work in yeast (**Fig. S3A-B**) and analysed protein-chromatin associations in small and large cells by mass spectrometry. The recruitment of all subunits of RNAPII to the genome increased in larger cells, as expected, and so did the initiation and elongation factors that regulate and associate with RNAPII (**Fig. 2B & S3E**). We note that only proteins first validated as being enriched on chromatin were analysed here to minimise potential confounding background effects (**Fig. S3C-D**). Taken together, these experiments show that more transcriptional machinery is expressed and loaded on the genome in larger cells, but does not delineate which, if any, component is dosage limiting in a manner that could couple global transcription to cell size.

**Figure 2.**
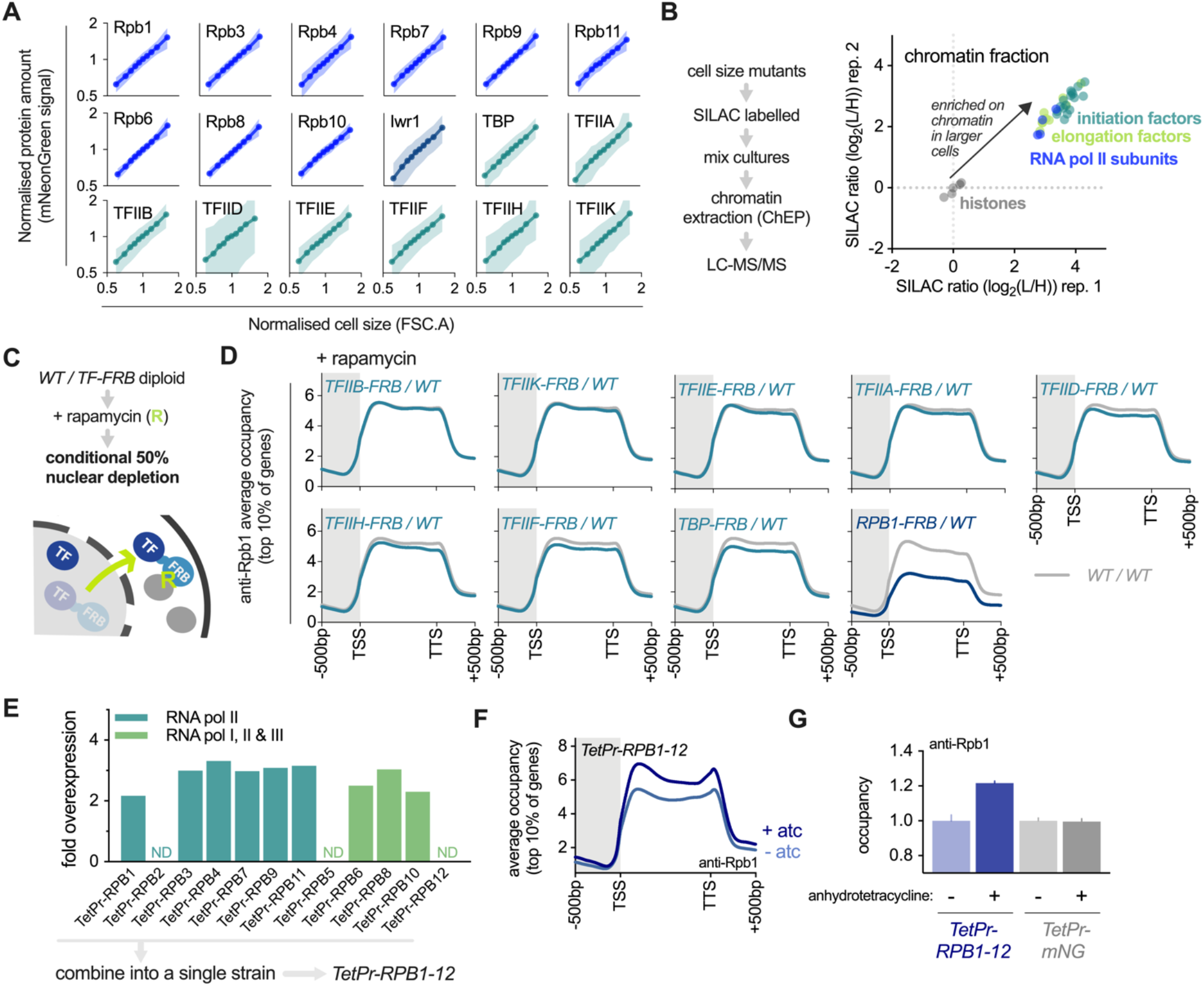
RNAPII is a major limiting component of the transcriptional machinery. See also Fig. S3-S5 **(A)** Protein amount (mNeonGreen signal) plotted against cell size (forward scatter) for subunits of the RNAPII pre-initiation complex determined by flow cytometry. The mean (±SD) is shown for each cell size bin. **(B)** Chromatin association of transcriptional machinery measured by mass spectrometry. SILAC-labelled cells of different sizes (Fig. S3E) were mixed and then chromatin was extracted (ChEP) and analysed by LC-MS/MS (see Fig. S3B for controls). Each axis shows an independent biological replicate for the normalized SILAC ratio of chromatin association between large (L) and small (H) cells. RNAPII subunits, initiation factors, and elongation factors are shown in comparison to histones. Only chromatin-enriched proteins are shown (Fig. S3C-D). **(C)** Schematic illustrating the 50% conditional depletion stratergy to test for dosage-limitation; a RNAPII pre-initiation complex subunit (TF) is tagged with the FRB domain (Haruki et al., 2008) and then crossed with a *WT* haploid to make a heterozygous diploid with one of the two alleles is FRB tagged, allowing for conditional depletion of half of that subunit from the nucleus. **(D)** Average Rpb1 occupancy across gene bodies of the top 10% of genes measured by spike-in normalized ChIP-seq in wild-type diploid cells (*WT/WT*) or diploids where 50% of the indicated factor was depleted from the nucleus upon rapamycin treatment. Rpb1-FRB anchor away results in efficient co-depletion of Rpb3 (Fig. S4A) suggesting that the whole RNAPII complex is efficiently co-depleted. Nuclear depletion is efficient and near-complete (i*.e.*, ∼50%) because ChIP against total Rpb1 and ChIP against a FLAG tagged non-depleted allele give a similar result (Fig. S4C-E). **(E)** The fold overexpression of individual RNAPII subunits from the TetPr after 45 minutes of anhydrotetracycline (atc) treatment. Expression levels from the TetPr was quantified using C-terminal mNeonGreen tagged proteins measured by flow-cytometry, which was then compared to the endogenously tagged allele of the respective subunit to calculate the fold overexpression (Fig. S5A). For *RPB2*, *RPB5* and *RPB12* this was not determined (N.D.) because tagging causes severe growth defects. All 12 subunits were then integrated into a single strain (see methods for details) to construct the *TetPr-RPB1-12* strain used in (F-G). 45 minute atc treatment results in approximately 2-3x overexpression of each subunit. **(F)** Average Rpb1 occupancy across the gene bodies of the top 10% of genes for the *TetPr-RPB1-12* strain before (-atc) or 45 minutes after (+atc) simultaneous overexpression of all 12 RNAPII subunits. **(G)** The global Rpb1 occupancy measured by spike-in normalized ChIP-seq after atc induced expression of either all RNAPII subunits (*TetPr-RPB1-12*) or free mNeonGreen (*TetPr-mNG*). Mean (±range) is plotted.

### RNAPII is a major limiting sub-complex of the PIC

To directly test potential limiting factors, we next performed a local perturbation to the nuclear amounts of each PIC component using the anchor away approach to conditionally deplete a targeted factor from the nucleus (Haruki et al., 2008). We constructed heterozygous diploids where one of the two alleles of a given PIC subunits is FRB tagged, allowing us to use rapamycin to rapidly and conditionally deplete half of it from the nucleus to a cytoplasm FKBP12 anchor (**Fig. 2C**). To determine how twofold depletion affects RNAPII loading, we then performed anti-Rpb1 spike-in normalized ChIP-seq. The general transcription factor complexes are not significantly limiting because a twofold reduction in their nuclear amounts results in at most a 5-10% reduction in global RNAPII occupancy (**Fig. 2D & S4**). However, depletion of half the RNAPII complex from the nucleus led to a much larger ∼40% reduction in occupancy (**Fig. 2D**), suggesting that RNAPII is a major limiting factor. Using a FLAG tag to quantify only the non-depleted allele in a *RPB1-FLAG/RPB1-FRB* strain, we confirmed the ∼40% value (**Fig. S4D-E**). Thus, our conditional depletion analysis shows RNAPII is a major limiting component of the PIC, while the other sub-complexes of the PIC are not.

### Transient overexpression is sufficient to increase RNAPII loading

If RNAPII is a limiting factor for its own recruitment to the genome, an increase in the amounts of the 12 subunit RNAPII complex should be sufficient to increase polymerase loading. To test this, we first optimised the overexpression conditions of individual subunits by comparing the overexpressed protein amounts (*e.g., Tetpr-RPB1-mNeonGreen*) with that of their endogenously expressed counterpart (*e.g., RPB1pr-RPB1-mNeonGreen*) (**Fig. S5**). This allowed us to define an induction regime in which all tested subunits are 2-3 fold overexpressed from an anhydrotetracycline-inducible promoter (*TetPr*) after a 45 minute induction (**Fig. 2E**). Next, we engineered a single yeast strain in which all 12 subunits can be simultaneously and conditionally overexpressed (*TetPr-RPB1-12*). In this strain, we observed a robust increase in global RNAPII occupancy across the genome after induction while there was no equivalent change in a control strain expressing mNeonGreen from the *TetPr* integrated at the same loci (**Fig. 2F-G**). Thus, RNAPII is a limiting component of the transcriptional machinery because RNAPII loading on the genome is sensitive to increases or decreases in its concentration.

### RNAPII occupancy increases non-linearly with size

Having established RNAPII as a major limiting component of the PIC we sought to examine more quantitatively how this could drive the scaling transcription with cell size. Prior theoretical models had proposed that a limiting factor(s) should be titrated against the genome (Lin and Amir, 2018; Sun et al., 2020). In these models, protein amounts - rather than concentrations - are critical because the on rate for binding is so high that nearly all molecules are associated with the genome. This is predicted to result in a directly proportional linear increase in DNA-binding with cell size.

To test this genome-titrated model, we first sought to simplify our experimental set up and measure RNAPII occupancy in cells of different sizes but with a fixed DNA amount. To do this, we isolated small G1 cells by centrifugal elutriation and arrested them in G1 by shutting off G1 cyclin expression. By arresting cells for increasing amounts of time, we generated populations with increasing cell sizes but with the same 1N DNA content (**Fig. 3A & S6A-C**). As expected, RNAPII occupancy increases with cell size but, crucially, this increase is sub-linear as it clearly deviates from a direct linear proportionality in larger cells (**Fig. 3B**). Importantly, these trends are not an artefact of elutriation (**Fig. S6E-F**) and are also clearly apparent in our measurements of asynchronously cycling cell size mutants (**Fig. 1E**).

**Figure 3.**
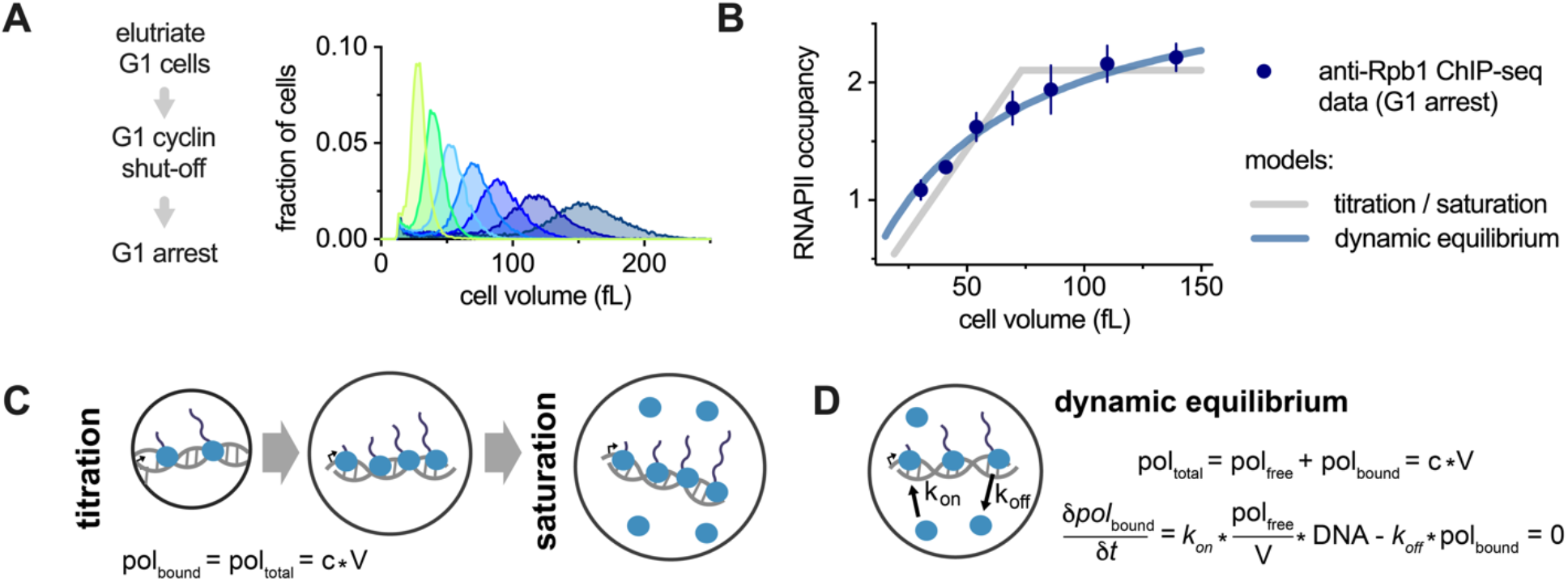
RNAPII loading is not directly proportional to cell size. See also Fig. S6 **(A)** Cell volume distributions determined by Coulter counter for G1 cells of different sizes that were used to analyse Rpb1 occupancy in (B). Small G1 cells were collected by centrifugal elutriation and then arrested in G1 using a conditional cyclin shut-off system to generate populations of G1 cells of increasing size, which were then collected for spike-in normalized ChIP-seq. See Fig. S6A for experimental design. **(B)** Rpb1 (anti-Rpb1) occupancy per cell plotted as a function of cell size in the indicated populations of G1 cells shown in (A). Each point shows the mean of two biological replicates (±range). The fit of two different mathematical models to these data are also show. (**C-D**) Schematics illustrating two alternative modes that can account for the data in B: (**C**) the previously proposed “titration / saturation” model, and (**D**) the “dynamic equilibrium” model proposed here.

The sub-linear increase in RNAPII occupancy with cell size rules out a simple titration model, so we considered two alternative possibilities. The first is a modification of the titration model in which, at a specific size threshold, the genome becomes saturated and transcription cannot increase further (Lin and Amir, 2018) (**Fig. 3C**). The second “dynamic equilibrium” model (**Fig. 3D**) assumes that the genome does not saturate and that the rate at which the free RNAPII, *pol_free_*, associates with the genome, *DNA*, is determined by mass action kinetics:

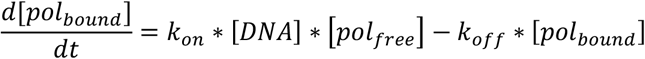

which can be rewritten in terms of amounts:

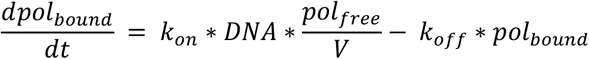

These kinetics are at steady state because the time scale of transcription is on the order of 1 minute, while the time scale of appreciable cell growth is on the order of 1 hour so that 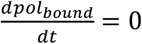. Finally, because the total RNAPII amount is proportional to cell size (*pol_total_* = *pol_bound_* + *pol_free_* = *c* ∗ *V*), where *c* is a constant, and the nuclear volume is always approximately 10% of cell volume (Jorgensen et al., 2007; Neumann and Nurse, 2007) we can solve for the amount of DNA bound RNAPII:

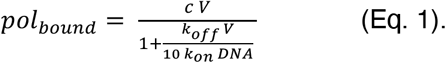

When fit to the data, the dynamic equilibrium model fits our global RNAPII occupancy measurements similarly well to the titration-saturation model (**Fig. 3B**). We note that the dynamic equilibrium model only has two parameters where, in essence, one is the concentration of RNAP II and the other is the dissociation constant for the interaction of RNAP II and the genome.

### RNAPII binding is driven by dynamic equilibrium kinetics

While both models can account for the amount of RNAPII bound, they make dramatically different predictions for what fraction of RNAPII is associated with the genome. The titration model predicts that nearly all nuclear RNAPII should be engaged on the genome resulting in a near-negligible free nucleoplasmic pool. As such, the genome-bound fraction should be fixed until cells surpass the saturation threshold at which point the bound fraction should decrease in proportion to the inverse of the cell volume 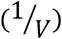. In contrast, the fit of the dynamic equilibrium model to our ChIP-seq data predicts that approximately 50% of nuclear RNAPII should be freely diffusing in the nucleoplasm of a 50 fL G1 cell, and that the bound fraction should decrease gradually and continuously as cell size increases so that: 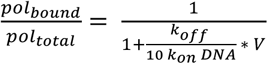.

To distinguish between these models, we sought to measure the fraction of bound and free RNAPII in single cells. We imaged and tracked single molecules of Rpb1-HALO and estimated their radius of gyration. Freely diffusing molecules diffuse rapidly across the nucleus and have a large radius of gyration, while chromatin-bound molecules are relatively immobile and have a small radius of gyration. Consistent with expectations, the majority of H2B histone molecules are tightly bound and the vast majority of nuclear mCitrine-NLS fluorescent molecules are highly mobile (**Fig. 4A-B**). Crucially, only half of the nuclear RNAPII subunit Rpb1 is freely diffusing, and the other half is bound (**Fig. 4B**). Moreover, the bound fraction of Rpb1 in individual G1 cells decreases gradually with cell size in broad agreement with the prediction of the dynamic equilibrium model (**Fig. 4C**). In contrast, these data show a very poor correspondence to the prediction of the titration-saturation model (**Fig. 4C**). We emphasize that the two free parameters for the equilibrium model were determined by fitting to the anti-Rpb1 ChIP-seq data (**Fig. 3B**) and are not adjusted in response to the single-molecule imaging data.

**Figure 4.**
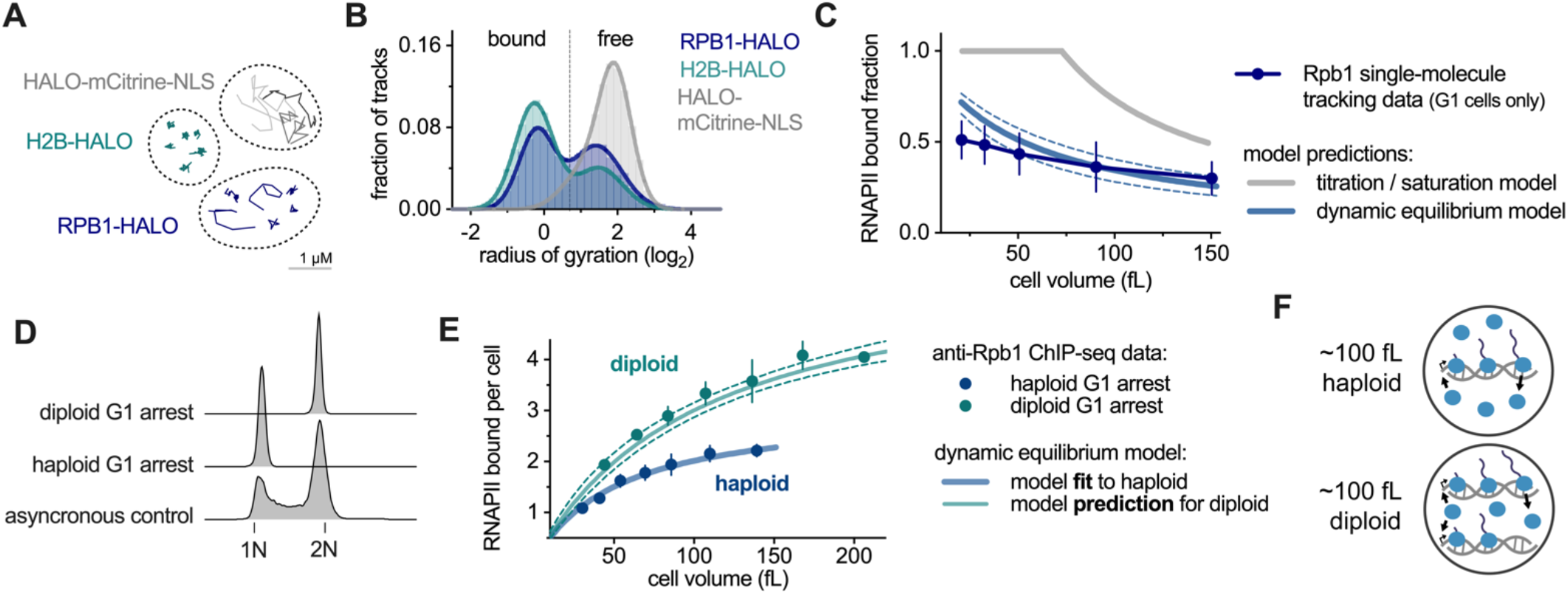
A dynamic equilibrium model predicts how cell size determines the recruitment of RNAPII to the genome. See also Fig. S6 **(A)** Example single molecule tracks in representative nuclei of the indicated genotype. **(B)** Histogram of the radius of gyration determined by single molecule imaging for HALO-mCitrine-NLS fusion proteins (n = 8057 tracks), Htb1 (n = 35978 tracks), and Rpb1 (n = 16008 tracks). The bimodal histogram of Rpb1 radii of gyration indicate chromatin-bound and freely diffusing fractions. **(C)** The fraction of bound Rpb1 molecules plotted as a function of cell size. Only G1 cells were analyzed. The mean for each cell size bin is plotted. Data shown are a combination of *WT* (n=182 cells) and *cln3Δ* (n=87 cells) cells. *cln3Δ* were used to increase the range of G1 cell sizes. **(D)** DNA content analysis by flow cytometry of haploid or diploid cells arrested on G1. **(E)** Rpb1 (anti-Rpb1) occupancy per cell plotted as a function of cell size in haploid and diploid populations of G1 arrested cells. Each point shows the mean of two biological replicates (±range). The dynamic equilibrium model prediction for diploids is calculated using the fit to the haploid data. 95% confidence intervals (dashed line) were generated using 10,000 bootstrap samples of the haploid data. **(F)** Summary schematic illustrating the effects of increase ploidy on DNA-bound RNAPII.

In our dynamic equilibrium model, the RNAPII loading rate is determined by the mass action kinetics of unbound polymerase and its target sites on the genome (**Fig. 3D**). As such, the amount of RNAPII on the genome should be sensitive to cell size *and* the amount of DNA. To test this, we determined the effects of increasing the genome copy number by performing the same elutriation experiment with diploid cells (**Fig. 4D** & **S6A-D**). These data show that RNAPII occupancy is significantly higher in diploids compared to haploid cells of the same size and show a very close correspondence to the model prediction for doubling DNA amount (**Fig**. **4E-F**). Importantly, this prediction is made using no additional free parameters as we simply double the value of ‘*DNA*’ in Eq. 1 above after fitting to the haploid data.

Taken together, our quantitative ChIP-seq and single-molecule tracking experiments are inconsistent with titration-based models. Instead, we propose that RNAPII occupancy is primarily determined via a surprisingly simple mass action dynamic equilibrium model. As a cell grows and synthesises more RNAPII, new RNAPII first enters the free fraction which increases the free nucleoplasmic concentration. This then drives more polymerase onto the genome until a new equilibrium is established at the increased size. However, this increase in the free concentration is not in direct proportion to cell size resulting in a sub-linear scaling of RNAPII occupancy on the genome that becomes more pronounced as cells get larger. This sub-linear increase is not due to saturation of the genome as previously thought. Instead, this is due to a progressively smaller fraction of polymerase being depleted from the nucleoplasm onto the genome in larger cells so that there is a smaller and smaller increase in the concentration of free nuclear polymerase that determines loading kinetics as cells enlarge.

### The chromatin template is invariant with cell size

A key component of our dynamic equilibrium model is that the on-rate (*k_on_*) is independent of cell size so that changes in the rate loading is only determined by changes in the concentration of free nucleoplasmic RNAPII. One prediction of this model is that chromatin should not become more or less permissive to polymerase recruitment at different cell sizes. To test this prediction that chromatin accessibility is size-independent, we adapted the dual-enzyme single-molecule footprinting (dSMF) assay (Krebs et al., 2017) for use in yeast. Briefly, this involves treating nuclei with a combination of CpG and GpC methyltransferases so that more accessible DNA is methylated, and protected DNA is not methylated. Differences in methylation patterns are then quantified by sequencing and normalized to a spike-in (**Fig. 5A**). When we performed dSMF on cells ranging from ∼35 fL to ∼100 fL we observed no major changes in chromatin accessibility between cells of different sizes (**Fig. 5B & S7A**). Consistent with this, we also do not see changes in histone occupancy measured by spike-in normalized ChIP-seq (**Fig. S6D**).

**Figure 5.**
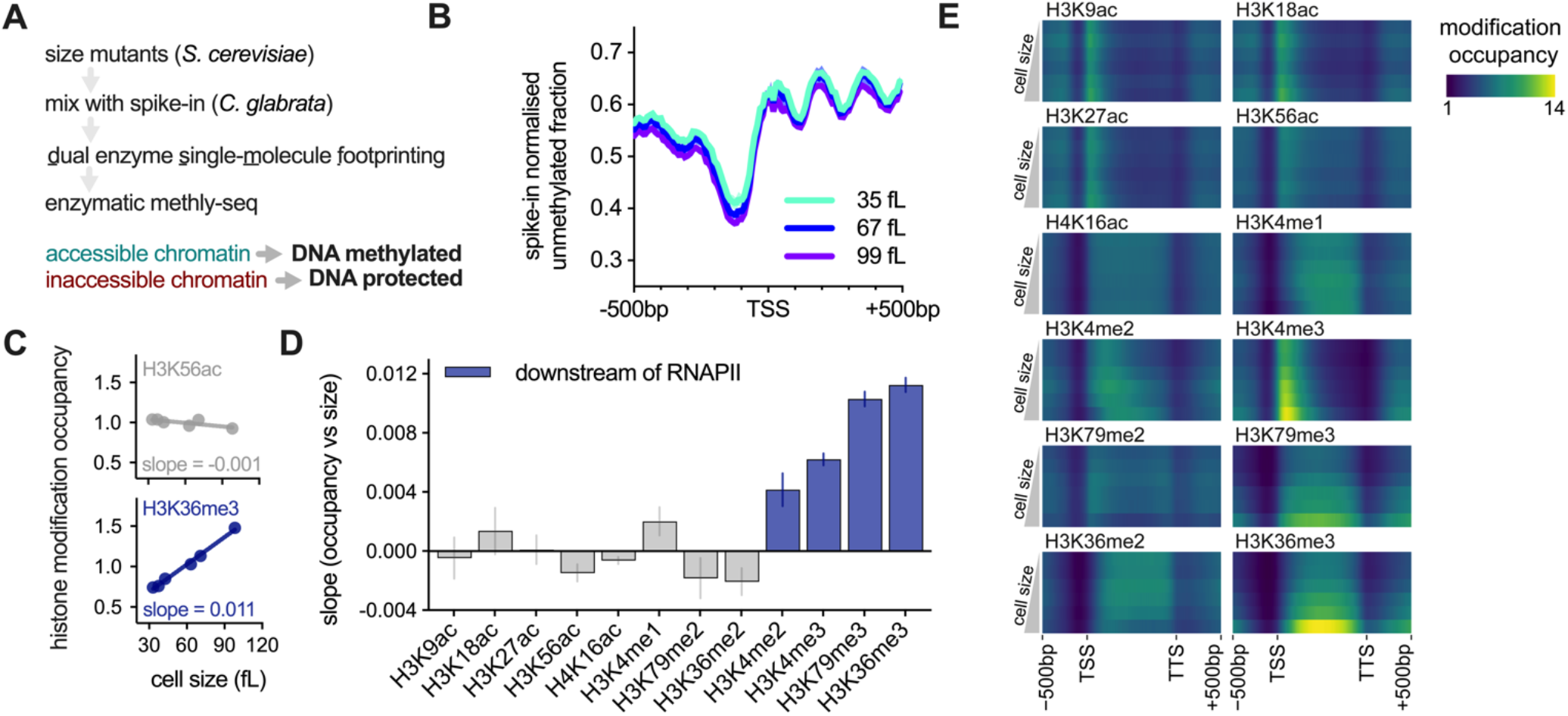
The upstream chromatin environment does not show major changes with cell size. See also Fig. S7 **(A)** Schematic of the workflow for measuring global chromatin accessibility in different cell size mutants using spike-in normalized <u>d</u>ual enzyme <u>s</u>ingle <u>m</u>olecule footprinting (dSMF). **(B)** Mean unmethylated DNA fraction plotted around transcriptional start sites (TSS) following dSMF in cell size mutants (low *WHI5* = 35 fL, medium *WHI5* = 67 fL, high *WHI5* = 99 fL; Fig. S7A). Unmethylated DNA corresponds to inaccessible chromatin and methylated DNA corresponds to accessible chromatin. Mean (±range) is plotted. (**C-E**) Histone modification occupancy measured in the different cell size mutants shown in Fig. 1D. **(C)** The occupancy for two example modifications (top: H3K56ac; bottom: H3K36me3) plotted against cell size. See also Fig. S7B. **(D)** The slope (±SE) for the linear fit between histone modification occupancy and cell size. See (C) for example fits for H3K56ac and H3K36me3. Larger slopes correspond to increased modification occupancy with cell size. Modifications shown in blue are reported to be deposited downstream of RNAPII initiation and/or elongation (see main text). **(E)** Average occupancy across gene bodies for the indicated modification. Each row corresponds to a different cell size mutant, ordered from small (top) to large (bottom) cell size.

Although the physical accessibility of chromatin is not changing with size, the chromatin landscape could still be modified via a changing pattern of histone modifications. To test this, we measured a panel of histone marks implicated in transcriptional processes by spike-in normalized ChIP-seq. The majority of modifications we measured did not change appreciably with cell size. However, four modifications associated with active transcription did increase significantly in larger cells: H3K4me2, H3K4me3, H3K79me3, H3K36me3 (**Fig. 5C-E & S7B**). Importantly, these di/tri-methyl modifications are known to be deposited downstream of transcriptional initiation or elongation, *i.e.*, their deposition depends on the recruitment of RNAPII (Krogan et al., 2003a; Krogan et al., 2003b; Ng et al., 2003; Santos-Rosa et al., 2002; Xiao et al., 2003). We therefore conclude that the chromatin landscape, both in terms of accessibility and histone modifications upstream of transcription, is largely invariant with cell size.

### mRNA decay feedback compensates for sub-linear transcriptional scaling

Our measurements and model for limiting RNAPII show that transcriptional scaling is not proportional to cell size, and instead displays a sub-linear trend that increasingly deviates from direct proportionality. This has two possible consequences: either mRNA amounts follow the same trends and mRNA concentrations decline in larger cells, or there is a buffering mechanism that stabilizes mRNA as size increases to compensate for the non-linear transcriptional scaling and maintain constant mRNA concentrations (**Fig. 6A**).

**Figure 6.**
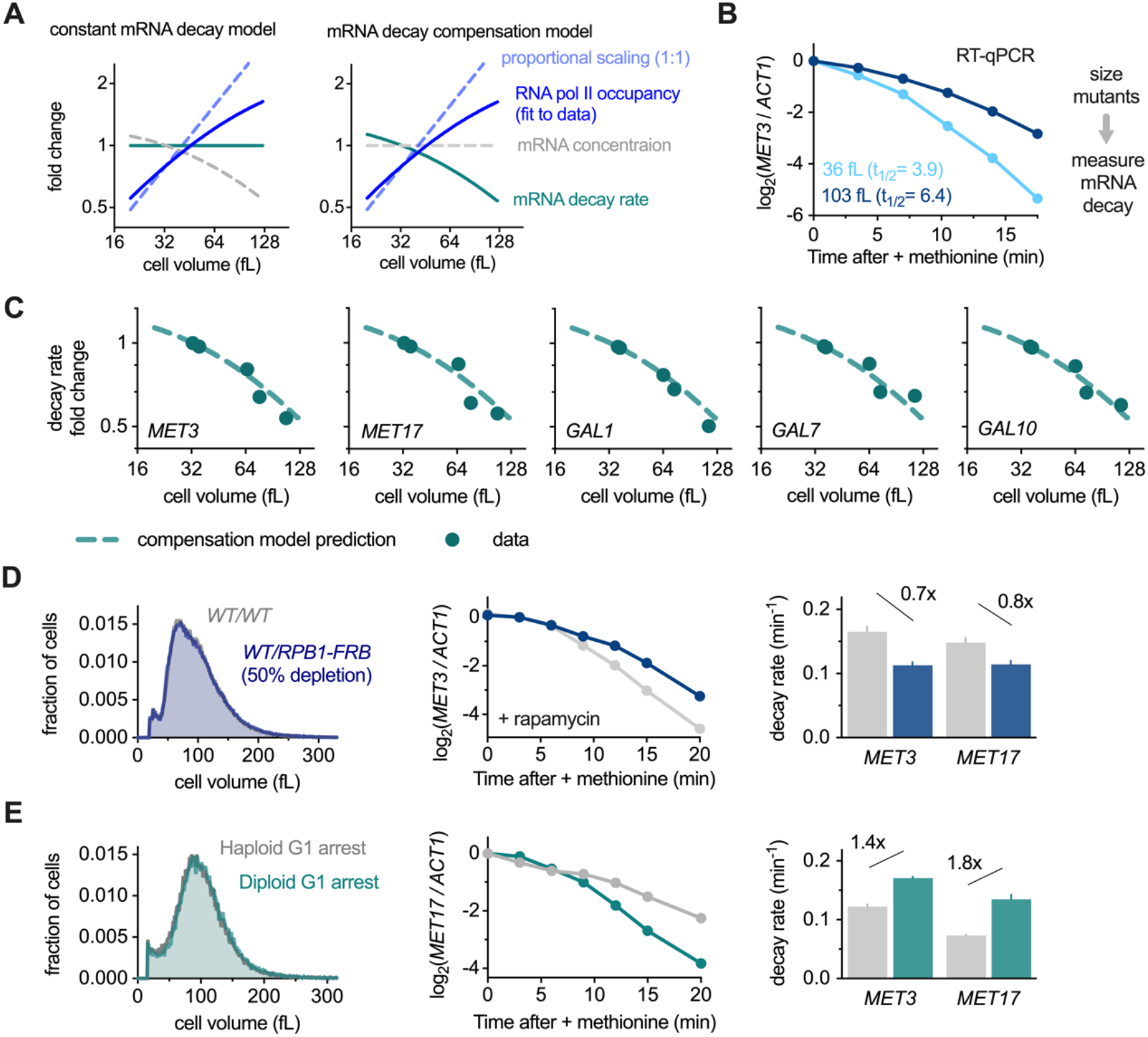
mRNA decay feedback compensates for the imperfect size-scaling of RNAPII activity. See also Fig. S8 **(A)** Predicted trends in global mRNA concentration and mRNA decay rate as a function of cell size that would result from the empirically observed transcription rate for either (left) no compensation or (right) feedback on mRNA decay rates that buffers mRNA concentrations as global RNAPII activity deviates from the proportional scaling trend. The empirical RNAPII occupancy trend is a fit of the dynamic equilibrium model (Fig. 3D) to the anti-Rpb1 ChIP-seq data of the cell size mutants presented in Fig. 1E. **(B)** *MET3* mRNA levels relative to *ACT1* mRNA after methionine addition in small (low *WHI5*) and large (high *WHI5*) cells. See also Fig.S8A-C. **(C)** mRNA decay fold change for *MET3*, *MET17, GAL1, GAL7* and *GAL10* following transcriptional shutoff in cells of different sizes (see methods for details). Mean of two biological replicates is shown. Dashed line indicates the predicted fold-change from the mRNA decay compensation model shown in (A). **(D)** Comparison between wild-type diploid cells (*WT/WT*) or diploids where 50% RNAPII is depleted from the nucleus (*WT/RPB1-FRB*) after 40 minutes rapamycin treatment. Cell size distributions (left) *MET3* mRNA levels relative to *ACT1* mRNA after methionine addition (middle) and *MET3* and *MET17* decay rates (right) are shown. Cell size distribution and decay rates are the average of three biological replicates. **(E)** As in D but comparison between diploid and haploid strains that have been arrested in G1, by G1 cyclin shutoff until they reach the same size (∼100 fL). Cell size distribution and decay rates are the average of two biological replicates.

To test for the presence of size-dependent changes in mRNA stability, we measured the levels of endogenous transcripts following transcriptional repression. We utilized two different treatments where either methionine or glucose was added to the media to shut-off the *MET* or *GAL* genes respectively and measured mRNA levels, following shut-off, in the different cell size mutants. This clearly shows that mRNA stability increases in larger cells (**Fig. 6B & S8B-D**). We then asked how this compared quantitatively to what would be required to maintain an invariant mRNA concentration, given the size-dependence of the RNAPII occupancy we have measured. Strikingly, the relative change in mRNA decay rate for all 5 transcripts that we examined aligned very closely to that predicted by a perfect mRNA decay compensation model (**Fig. 6C**). Thus, mRNA decay rates are precisely adjusted to compensate for the imperfect scaling of RNAPII activity with size to ultimately ensure mRNA concentrations are kept constant as cell size increases.

Although we do not know the precise mechanism for this compensation, we believe that it is due to feedback on global mRNA concentrations, rather than a direct coupling to cell size. This is because we observed a similar decrease in mRNA turnover rates in cells subject to a rapid depletion of half the nuclear RNAPII, even though cell size does not change in this context (**Fig. 6D & S8E**). Similarly, we observed that ∼100 fL G1 haploid cells have lower mRNA turnover rates than similarly sized diploids (**Fig. 6D & S8E**) consentient with the lower global transcription rate in haploids of this cell size (**Fig. 4E-F**).

## DISCUSSION

How cells scale global mRNA amounts to remain in proportion to cell size during growth has been a long-standing and fundamental question in cell biology (Elliott, 1983a, b; Elliott and McLaughlin, 1979; Fraser and Nurse, 1978, 1979; Padovan-Merhar et al., 2015; Sun et al., 2020; Zhurinsky et al., 2010). It was previously suggested that the size-scaling of mRNA was due to mRNA synthesis rates increasing in direct proportion to cell size. In these models, such size-scaling is due to an unknown limiting transcription factor that is titrated against the genome (Lin and Amir, 2018; Sun et al., 2020).

### Transcriptional scaling via a dynamic equilibrium model for limiting RNAPII

Here, we empirically identified RNA polymerase II (RNAPII) as a major limiting factor by showing that polymerase loading onto the genome is sensitive to both acute increases and decreases in its own concentration, whereas other components of the transcriptional initiation machinery are not limiting (**Fig. 2**). This finding is especially satisfying given, to our knowledge, it was first speculated that RNA polymerase could be the hypothetical limiting factor four decades ago (Elliott, 1983b). However, our single-molecule imaging reveals that RNAPII is not titrated against the genome as there is a significant pool of RNAPII that is freely diffusing in the nucleoplasm. Moreover, the amount of active RNAPII does not scale in direct proportion with cell size so that larger cells exhibited systematically larger deviations from the amount RNAPII engaged on the genome predicted by the titrated factor model. Instead, we propose a new dynamic equilibrium model in which the loading of RNAPII on the genome is determined by mass action kinetics with a defined on-rate *k_on_*_’_, which is independent of cell size (**Fig. 3-4**). The size-independence of *k_on_* is consistent with our measurements showing that there are no major changes in chromatin accessibility as cells grow larger (**Fig. 5**). In this model, as a cell grows, newly synthesised RNAPII increases the free nucleoplasmic concentration driving more polymerase onto the genome thereby establishing a new equilibrium between the bound and unbound RNAPII populations at the increased size. However, this increase is not in direct proportion to cell size, resulting in a deviation from linear scaling that becomes increasingly pronounced as cells get larger and transcription approaches a plateau as a function of cell size (**Fig. 7**). Together, our model quantitively predicts the relationships between RNAPII occupancy, cell size, and ploidy with only two free parameters. This model also accurately predicts the fraction of RNAPII that is bound to the genome and that which is freely diffusing in the nucleoplasm (**Fig. 3-4**).

**Figure 7.**
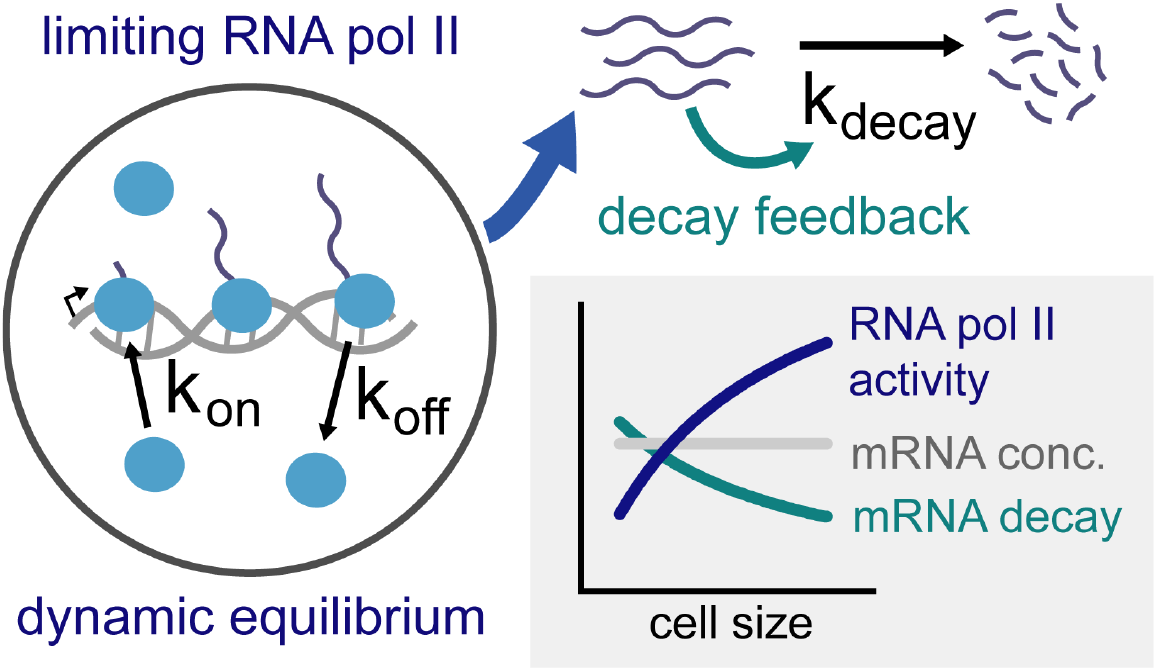
Summary schematic. A dynamic equilibrium drives more limiting RNAPII onto the genome in larger cells, resulting in a sub-linear increase in transcription with cell size. An additional feedback mechanism on mRNA decay rates increases mRNA stability in larger cells to compensates for the sub-linear transcriptional scaling and keep mRNA concentrations approximately constant as cells grow within the physiological size range.

### mRNA size-scaling requires additional buffering via feedback on mRNA turnover

While our dynamic equilibrium model successfully accounts for the size-dependent increase in RNAPII loading that we observe, it raises the question as to how the global mRNA concentration can be constant when transcription scales sub-linearly with cell size. If mRNA half-lives were independent of cell size as previously thought (Padovan-Merhar et al., 2015; Zhurinsky et al., 2010), mRNA concentrations would decrease in larger cells. Instead, we observe that mRNA stability decreases in larger cells to exactly compensate for the sublinear size-dependence of transcription (**Fig. 6**). Taken together, our data show how the mass action recruitment kinetics of limiting RNAPII and feedback on mRNA stability act as two mechanisms working in concert to ensure the precise scaling of mRNA with cell size (**Fig. 7**).

The presence of feedback operating on mRNA stability is highly reminiscent of the previous observations in both yeast and human cells that mutations affecting transcription are buffered by a compensatory decrease in mRNA turnover (Baptista et al., 2017; Berry et al., 2021; Helenius et al., 2011; Plaschka et al., 2015; Rodriguez-Molina et al., 2016; Schulz et al., 2014; Slobodin et al., 2020; Sun et al., 2012; Warfield et al., 2017). This is most dramatically seen in a recent study that examined global transcription rates and RNA concentrations in a genome-wide siRNA screen. While perturbations affecting transcription were numerous, these did not result in changes to RNA concentration (Berry et al., 2021). Thus, both growing larger and perturbation to transcriptional activity elicit a similar feedback effect to buffer mRNA concentrations. Consistent with this, we observed that yeast cells of the same size with quantitative differences in global RNAPII occupancy per cell also show corresponding changes in mRNA stability supporting the notion the size-dependent changes in transcript stability we observe are not due to a direct coupling to size. Thus, we anticipate that the feedback that maintains mRNA concentrations in growing cells and in response to transcriptional mutations arises from the same underlying molecular mechanism. Indeed, natural variations in cell size may provide the physiological context for which this mRNA stability feedback first evolved.

### Transcriptional scaling and optimal size ranges for cellular fitness and function

While the mechanisms we identify here sustain mRNA concentrations across a range of physiological sizes, it has also been shown that the scaling of macromolecule amounts with cell size ultimately breaks down when cells grow very large – *i.e.*, beyond the size range we examine here. This causes global RNA and protein concentrations to decrease above an upper cell size limit, effectively diluting the cytoplasm (Neurohr et al., 2019). This upper size limit is associated with a decline in many aspects of cellular physiology including attenuated signalling transduction, failure to re-enter the cell cycle, and defective conditional gene expression programs (Neurohr et al., 2019). Thus, for a given cell type there is an optimal size range, beyond which cells cannot grow efficiently and cellular processes stop working optimally.

The transition between optimal and non-optimal cell sizes is now linked to a range of physiological and pathological situations. For example, proliferation is attenuated by increased cell size, which has recently been isolated as a causal contributor to senescence – the tumour suppressive state of irreversible cell cycle exit characterised by massive cell enlargement (Cheng et al., 2021; Demidenko and Blagosklonny, 2008; Lanz et al., 2021; Neurohr et al., 2019; Wilson et al., 2021). Moreover, stem cells are amongst the tiniest in the body but tend to enlarge as an organism ages (Lengefeld et al., 2021; Li et al., 2015; Yang et al., 2018) and recent work has demonstrated that naturally or artificially enlarged haemopoietic or intestinal stem cells accelerate the ageing-associated decline in stem-cell function (Lengefeld et al., 2021). There is also a broad relevance to developmental biology as cells tend to enlarge as they differentiate, and the fact that cell types in the body vary over multiple degrees of magnitude suggests that each cell type exists at its own carefully optimised size.

The size-dependent patterns of global mRNA and protein scaling is likely to be critical for determining the optimal and non-optimal size range in some, and perhaps most, of these circumstances. However, it has not established why cells cannot scale biosynthesis with cell size above this upper size limit. Our data, in part, explains this because we observe that as cells enlarge RNAPII recruitment to genome does not increase in direct proportion to cell size and eventually reaches a plateau (**Fig. 1 & 3**). The plateau does not arise due to the saturation of the genome with RNAPII as has been previously proposed, but instead is a natural consequence of the mass action chemical kinetics we found to govern RNAPII loading. As cells grow larger the divergence between cell size and global transcription, which is initially small, becomes progressively more pronounced. Initially, global mRNA concentrations are not affected because the sub-linear transcriptional scaling is precisely buffered by feedback on mRNA decay rates to stabilise transcripts in larger cells. However, this balancing act between global transcription and decay eventually breaks down as the feedback on mRNA stability is no longer able to compensate for the ever-increasing divergence between transcription and cell size. Ultimately, this then results in the dilution of the global mRNA concentration.

Thus, we speculate that upper cell size limit corresponds to the point at which the sub-linear increase in transcription with cell size can no longer support efficient cellular biosynthesis. Consistent with this, doubling the ploidy increases the size-dependent loading of RNAPII onto the genome (**Fig. 4 D-F**) and also increases the upper cell size at which cells can function and grow efficiently (Neurohr et al., 2019). This provides a simple rationalisation as to why polyploidy is commonly coupled to increased cell size in nature – namely, polyploidy is a simple route to promote efficient cellular biosynthesis in bigger cells.

## SUPLEMENTAL FIGURES

**Figure S1.**
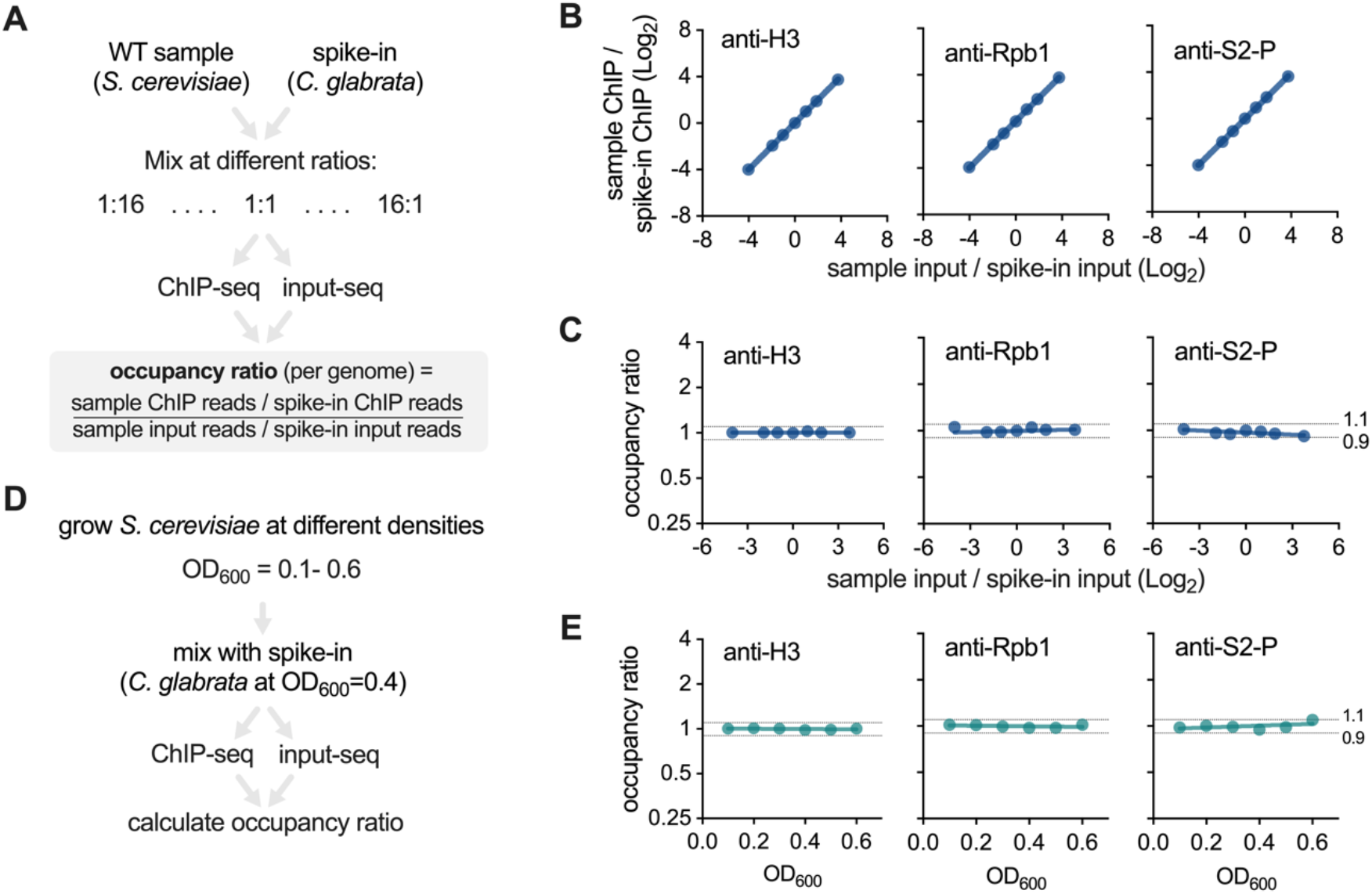
Controls for spike-in normalized ChIP-seq. Related to Fig. 1D **(A-C)** Mixing-ratio controls for spike-in normalized ChIP-seq. The design of the mixing-ratio control experiment is shown in **(A)**: *S. cerevisiae* and *C. glabrata* samples were mixed in 7 different ratios before anti-Rpb1, anti-S2P or anti-H3 ChIP-seq. Input samples were also sequenced to determine the mixing ratio. This experiment demonstrates the good dynamic range and linearity of spike-in normalized ChIP-seq because differences in the input ratio of *S. cerevisiae* to *C. glabrata* are reflected in proportional changes in the *S. cerevisiae* to *C. glabrata* ChIP ratio **(B)** and in no changes to the input normalized occupancy ratio **(C)**. ChIP and occupancy ratios are expressed the fold-change relative to the middle sample. This control is conceptually the same as to the reported for Scc1 ChIP by Hu *et al*. 2015 (Hu et al., 2015). This control also demonstrates that different *S. cerevisiae* samples do not need to be mixed at identical ratios with spike-in cells because variations in mixing ratio are linearly accounted for in the input normalized occupancy ratio. **(D-E)** Cell density control for spike-in normalized ChIP-seq. The design of the cell density control experiment is shown in **(D)**: *S. cerevisiae* cells were grown to different cell densities (OD600) mixed with approximately the same cell number of *C. glabrata*. The sample was then fixed and processed for spike-in normalized ChIP-seq. The occupancy ratio for anti-Rpb1, anti-S2P, or anti-H3 is constant as a function of cell densities **(E)** showing that *S. cerevisiae* samples do not need to be collected at identical cell densities to compare samples. All samples in the rest of the study were collected between OD600 = 0.2 and OD600 = 0.4 unless stated otherwise.

**Figure S2.**
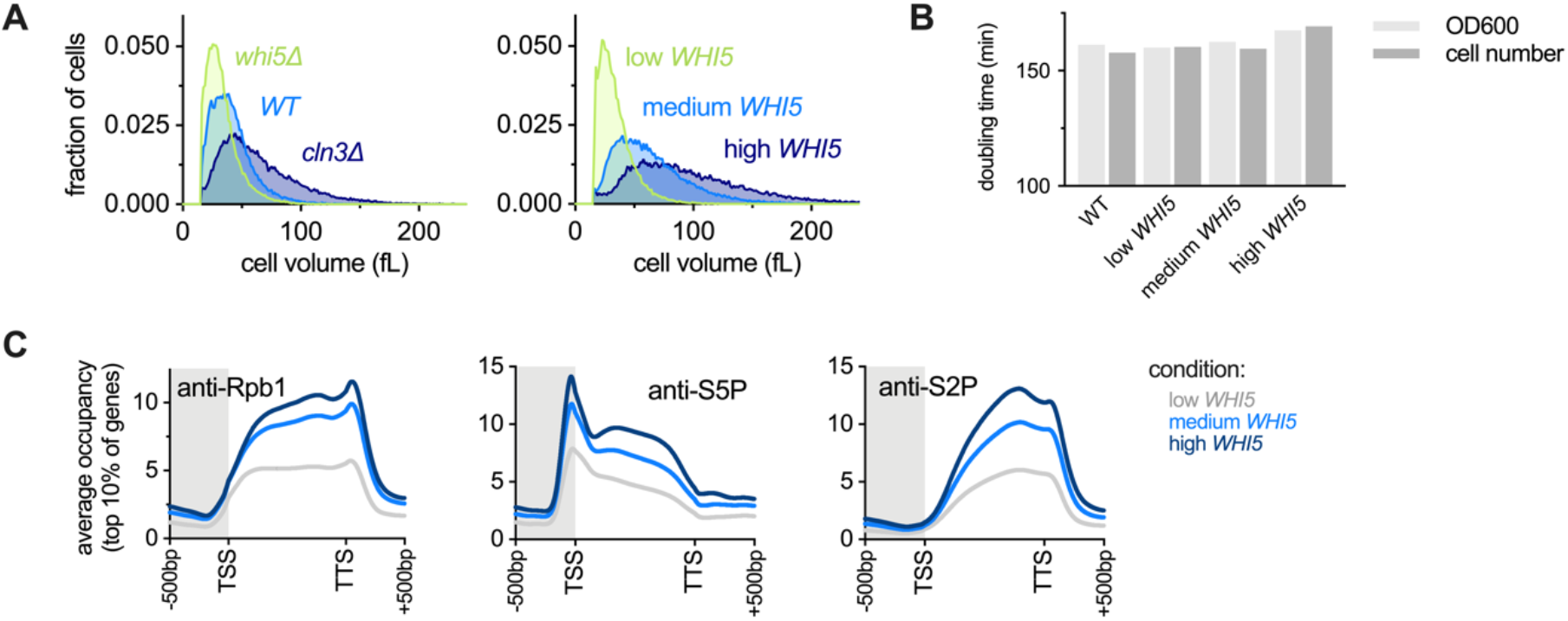
Cell size mutant Rpb1 spike-in normalized ChIP-seq supporting data. Related to Fig. 1C-F **(A)** Cell size distributions determined by Coulter counter for the cell size mutants used for the experiment in Fig. 1D-F. Expression of low, medium, and high *WHI5* expression was induced with 0mM, 10mM, and 30mM beta-estradiol, respectively. **(B)** Doubling time calculated from either OD600 or cell number measurements for cells expressing low, medium, and high levels of *WHI5* induced from a beta-estradiol responsive promoter with 0mM, 10mM, and 30mM beta-estradiol, respectively. Cultures were grown to steady-state before the growth curve measurements were started. **(C)** Average occupancy across the gene bodies of the top 10% of genes for total Rpb1, initiated Rpb1 (anti-S5-P), and elongating Rpb1 (anti-S2-P) in cells expressing low, medium, and high levels of *WHI5*. Global occupancy measurements for the same data are shown in Fig.1B.

**Figure S3.**
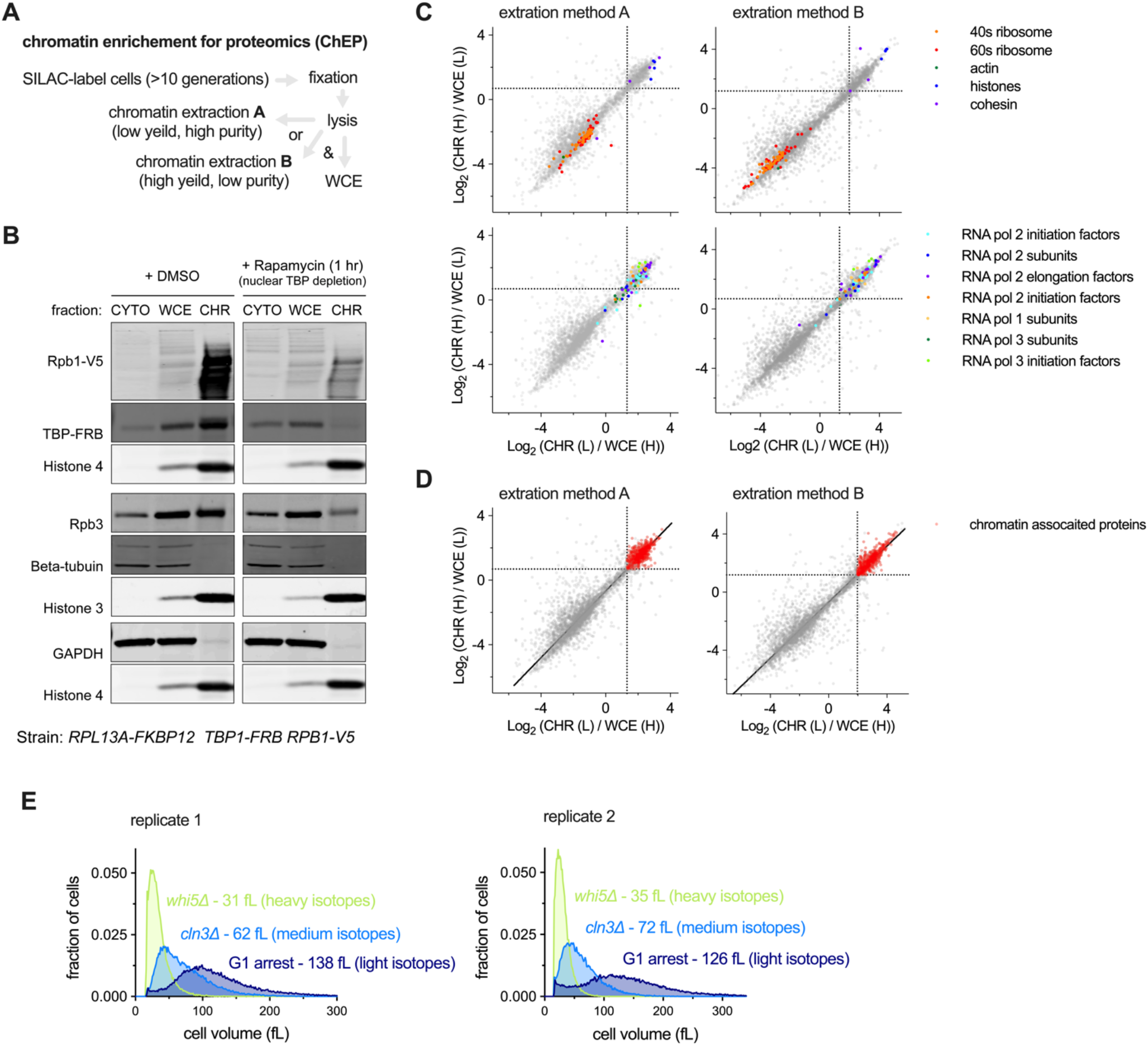
<u>Ch</u>romatin <u>e</u>nrichment for <u>p</u>roteomics (ChEP) workflow and controls. Related to Fig. 2B **(A)** Schematic of the workflow for chromatin extraction by ChEP. Both the high purity / low yield extraction (method A), and the low purity / high yield extraction (method B) were used. See methods for details. **(B)** Control experiment showing that ChEP can quantify changes in protein chromatin association. To test this, TPB and RNAPII chromatin association was conditionally prevented using a TBP-FRB strain where TBP is conditionally depleted from the nucleus upon rapamycin treatment thus preventing RNAPII recruitment to the genome. Upon rapamycin treatment for 1 hour, TBP and RNAPII (Rpb1 and Rpb3) have reduced chromatin association measured by ChEP compared to the DMSO control treatment. Thus, ChEP can be used to quantify changes in chromatin association. We note that, under these conditions, RNAPII is still present in the nucleus but is not associated with chromatin (See Fig. S4A), indicating that ChEP is unlikely to be subject to major background due to non-specific chromatin interactions. **(C-D)** Experiment to identify proteins enriched by ChEP. Cells were SILAC label with heavy (H) or light (L) isotopes and three extractions were recovered from each: whole cell extract (WCE), chromatin extraction A, and chromatin extraction B. WCE and chromatin with opposite SILAC labels where mixed and analysed by LC-MS/MS. Each axis shows an independent biological replicate. Left hand panels show chromatin extraction A (high purity, low yield). Right hand panels show chromatin extraction B (low purity, high yield). Known cytoplasmic factors (ribosomes and actin) and chromatin-associated proteins (RNA polymerase subunits, transcription factors, histones and cohesion) are shown in **(C)**, validating the chromatin extractions do enrich chromatin-associated factors. Proteins defined as chromatin associated are shown in red in **(D)**. A robust regression line is in black. Chromatin associated proteins were defined as proteins enriched in the chromatin fraction of both replicates for both extraction method A and extraction method B. The thresholds used to define chromatin-associated proteins are shown as dashed black lines (see methods for details). **(E)** Cell size distributions determined by Coulter counter for the SILAC labelled cultures used in the experiment in Fig. 2B. All strains have a SILAC-compatible genetic background (see methods for details).

**Figure S4.**
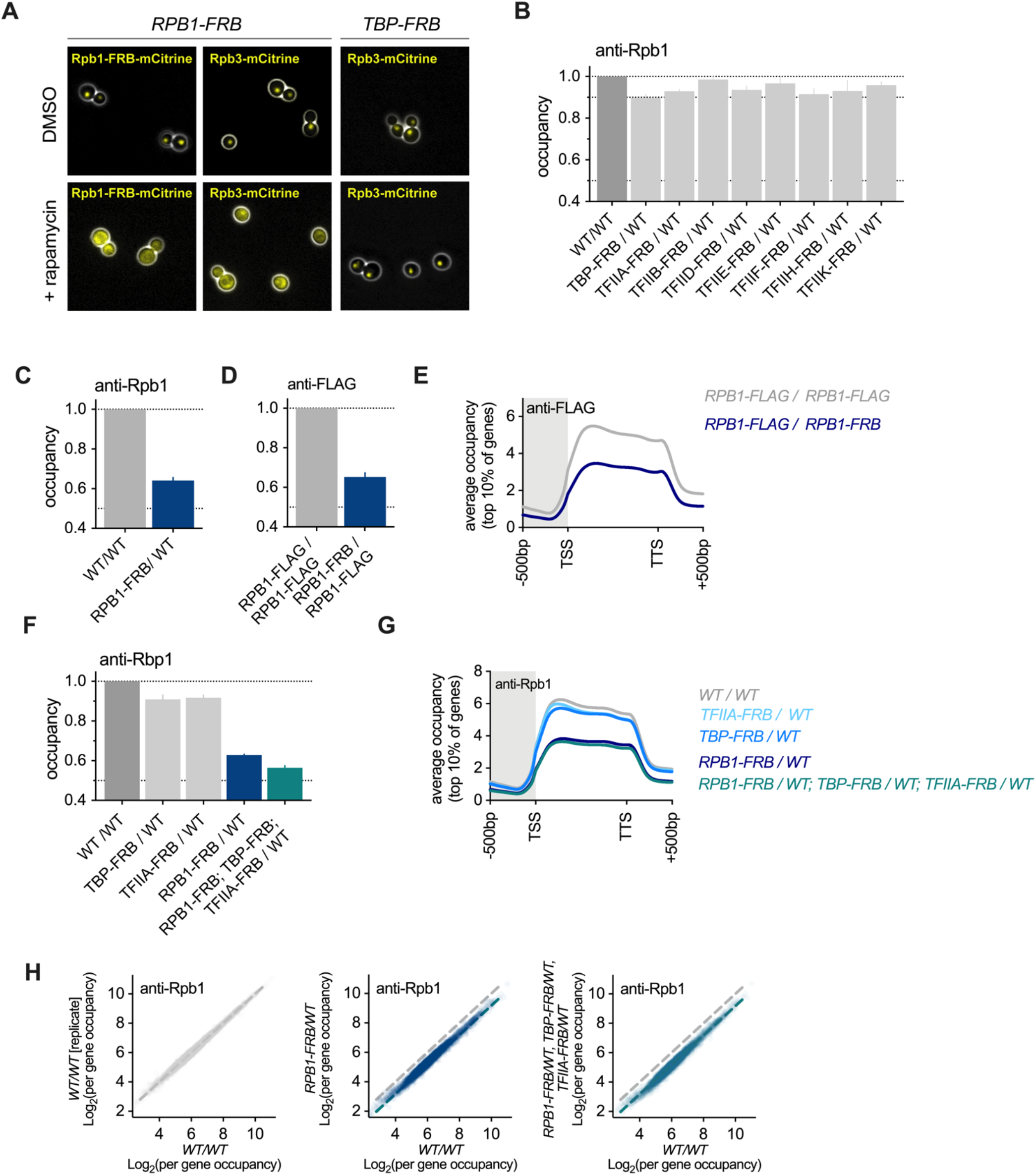
RNAPII 50% depletion and over-expression experiments supporting data. Related to Fig. 2C-D **(A)** Microscopy images (phase contrast in white, mCitrine signal in yellow) of the indicated genotypes treated with DMSO or rapamycin for 1 generation before imaging. Treatment of Rpb1-FRB cells leads to nuclear depletion of Rpb3 indicating the whole RNAPII complex is likely to be efficiently co-depleted. Treatment of TBP-FRB cells does not alter Rpb3 localization, indicating that different sub-complexes of the Pre-Initiation-Complex do not appear to co-deplete one another. **(B)** The global Rpb1 occupancy measured by spike-in normalized ChIP-seq in wild-type diploid cells (*WT/WT*) or diploids where 50% of the indicated RNAPII Pre-Initiation Complex subunit was depleted from the nucleus following a 100 minute rapamycin treatment. Mean (±SEM) is plotted. The corresponding average Rpb1 occupancy across the gene bodies of the top 10% of genes in these samples is shown in Fig. 2D. **(C)** The global Rpb1 occupancy measured by spike-in normalized ChIP-seq in wild-type diploid cells (*WT/WT*) or diploids where 50% of RNAPII was depleted from the nucleus by a 40 minute rapamycin treatment. Mean (±SEM) is plotted. The corresponding average Rpb1 occupancy across the gene bodies of the top 10% of genes in these samples is shown in Fig. 2D. **(D)** The global Rpb1-FLAG occupancy measured by spike-in normalized anti-FLAG ChIP-seq in wild-type diploid cells (*RPB1-FLAG/RPB1-FLAG*) or diploids where one allele is FLAG tagged and the other is depleted from the nucleus upon rapamycin treatment (*RPB1-FRB/RPB1-FLAG*). Mean (±SEM) is plotted. These data indicate that the nuclear depletion of Rpb1 is efficient and near-complete (i*.e.*, ∼50%) because ChIP against total Rpb1 in (C) and ChIP against the non-depleted allele in (D) give similar values. **(E)** The average Rpb1 occupancy across the gene bodies of the top 10% of genes for the samples shown in (D). **(F)** The global Rpb1 occupancy measured by spike-in normalized ChIP-seq in wild-type diploid cells (*WT/WT*) or diploids where 50% of the indicated RNAPII Pre-Initiation Complex subunits were depleted from the nucleus by a 40 minute rapamycin treatment. Data are also show in an experiment with the simultaneous 50% nuclear depletion TBP, TFIIA, and RNAPII. Mean (±SEM) is plotted. **(G)** The average Rpb1 occupancy across the gene bodies of the top 10% of genes for the samples shown in (F). **(H)** The mean per gene Rpb1 occupancy measured (left) in wild-type cells, (center) in cells after 50% nuclear depletion of RNAPII, or (right) in cells after the simultaneous 50% nuclear depletion TBP, TFIIA, and RNAPII. Each point corresponds to a single gene and shows the Rpb1 occupancy between TSS and TTS. Dashed lines show the average trends for the respective samples. Global occupancy values for the same samples are shown in (F).

**Figure S5.**
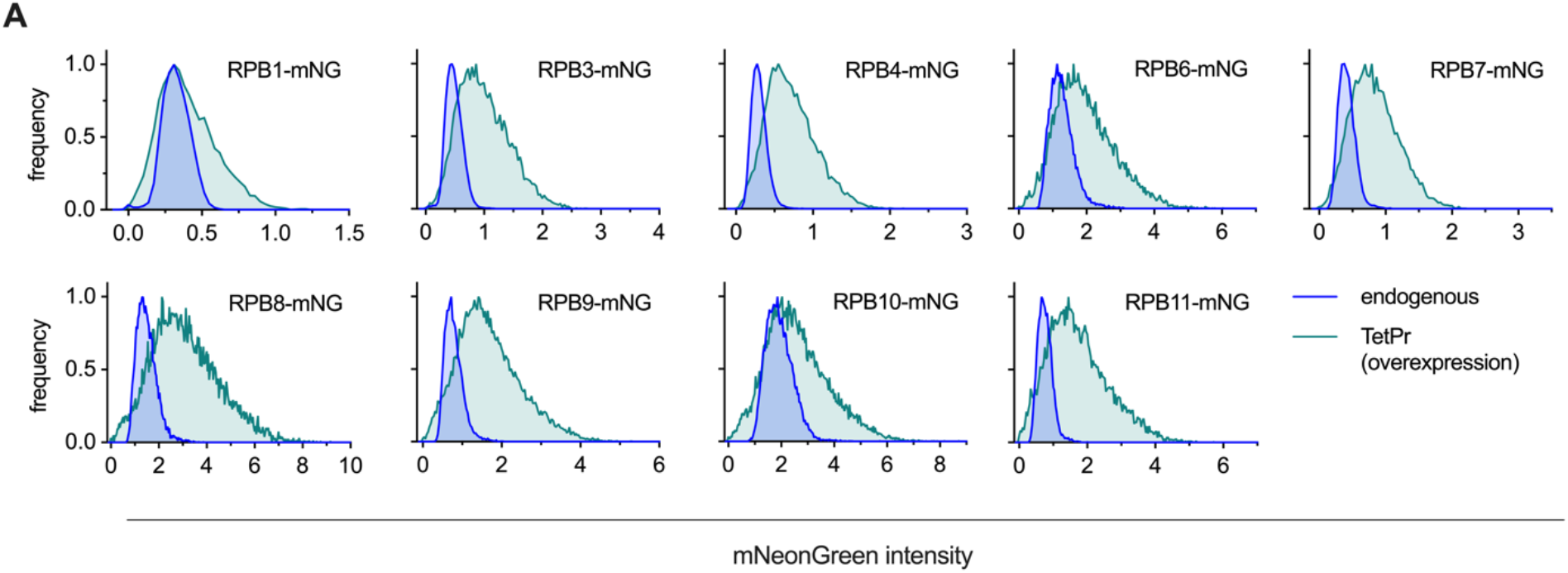
RNAPII over-expression experiments supporting data. Related to Fig. 2E-G **(A)** Histogram (mode normalized) of the mNeonGreen signal measured by flow-cytometry for the indicated RNAPII subunit C-terminal fusion with mNeonGreen (mNG). Fusion proteins were either expressed from their endogenous promoter (blue; endogenous) or overexpressed from a *Tet* promoter alongside the untagged endogenous copy of the same gene (green; TetPr). Expression from the TetPr was induced by anhydrotetracycline treatment (45 minutes). The fold overexpression that is induced upon *Tet* promoter expression was calculated by comparing the median intensity of the overexpressed construct to the endogenous allele and is shown in Fig. 2E.

**Figure S6.**
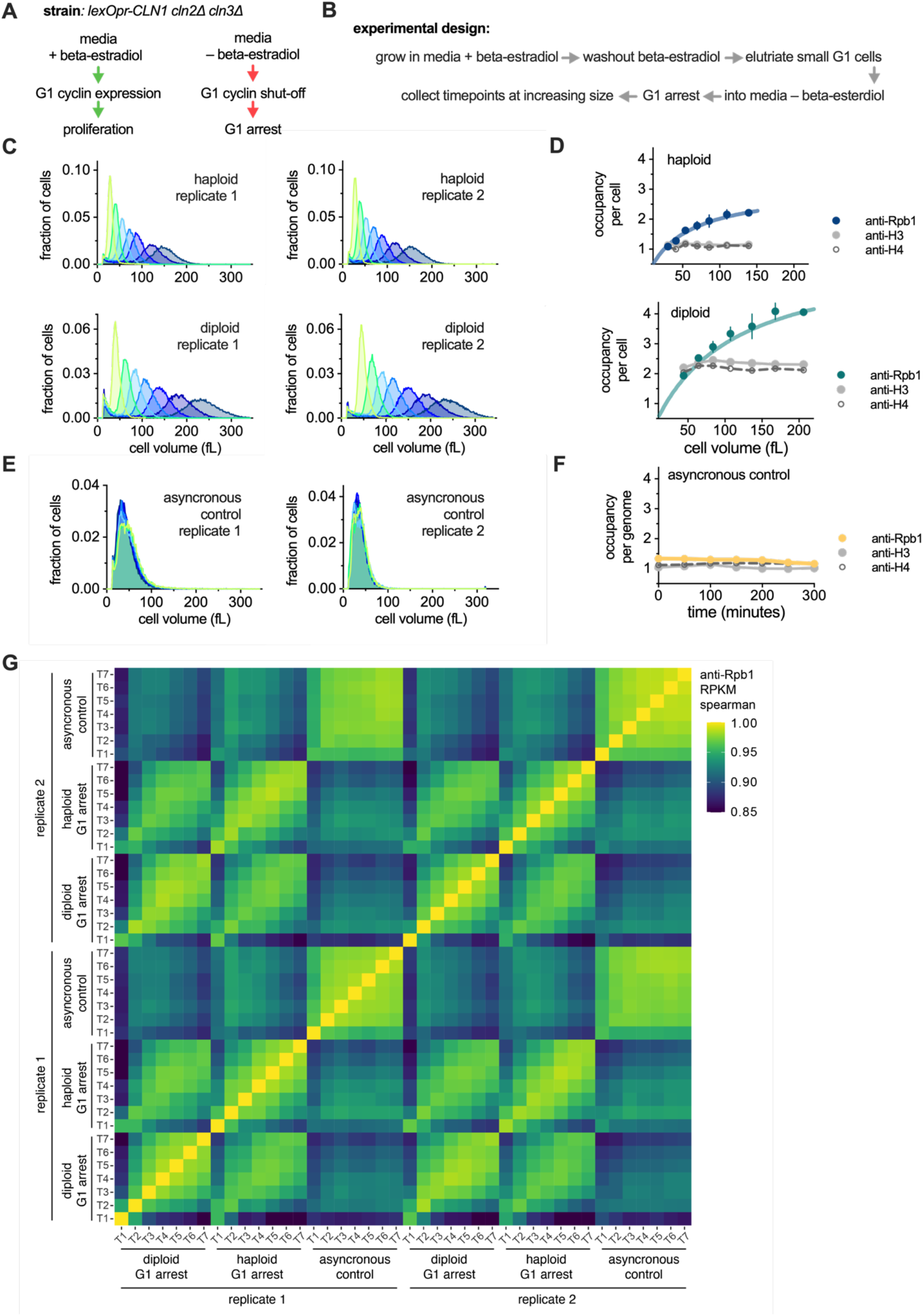
G1 arrest experiment supporting data and additional controls. Related to Fig. 3-4 **(A-B)** Schematic of the experimental design to generate populations of G1-arrested cells of increasing cell size. **(A)** A G1-cyclin shut-off strain was used that allows the conditional inactivation of G1 cyclin expression upon removal of beta-estradiol from the media. **(B)** Small G1 cells where then collected by centrifugal elutriation and populations of increasing cell size were generated as G1 arrest duration increases. See methods for details. **(C)** Cell size distributions determined by Coulter counter for G1 arrested haploid (top) and diploid (bottom) cultures of increasing cell size used to analyse Rpb1, Histone 3 and Histone 4 occupancy in (D) and Fig. 3-4. **(D)** Rpb1 (anti-Rpb1), Histone 3 (anti-H3) and Histone 4 (anti-H4) occupancy per cell plotted as a function of cell size in G1 arrested haploids (top) or diploid (bottom). The mean (±range) of two biological replicates is shown for anti-Rpb1, whereas only a single replicate of anti-H3 and anti-H4 is shown. The anti-Rpb1 data are also plotted in Fig. 3B and 4E. **(E)** Cell size distributions determined by Coulter counter for asynchronous control cultures used to analyse Rpb1, Histone 3 and Histone 4 occupancy in (F). **(F)** Rpb1 (anti-Rpb1), Histone 3 (anti-H3) and Histone 4 (anti-H4) occupancy per cell plotted as a function of cell size in asynchronous control cultures. The mean (±range) of two biological replicates is shown for anti-Rpb1 whereas only a single replicate of anti-H3 and anti-H4 is shown. **(G)** Heatmap showing the spearman rank correlation coefficient between the anti-Rpb1 RPKM values of all samples shown in (C-F).

**Figure S7.**
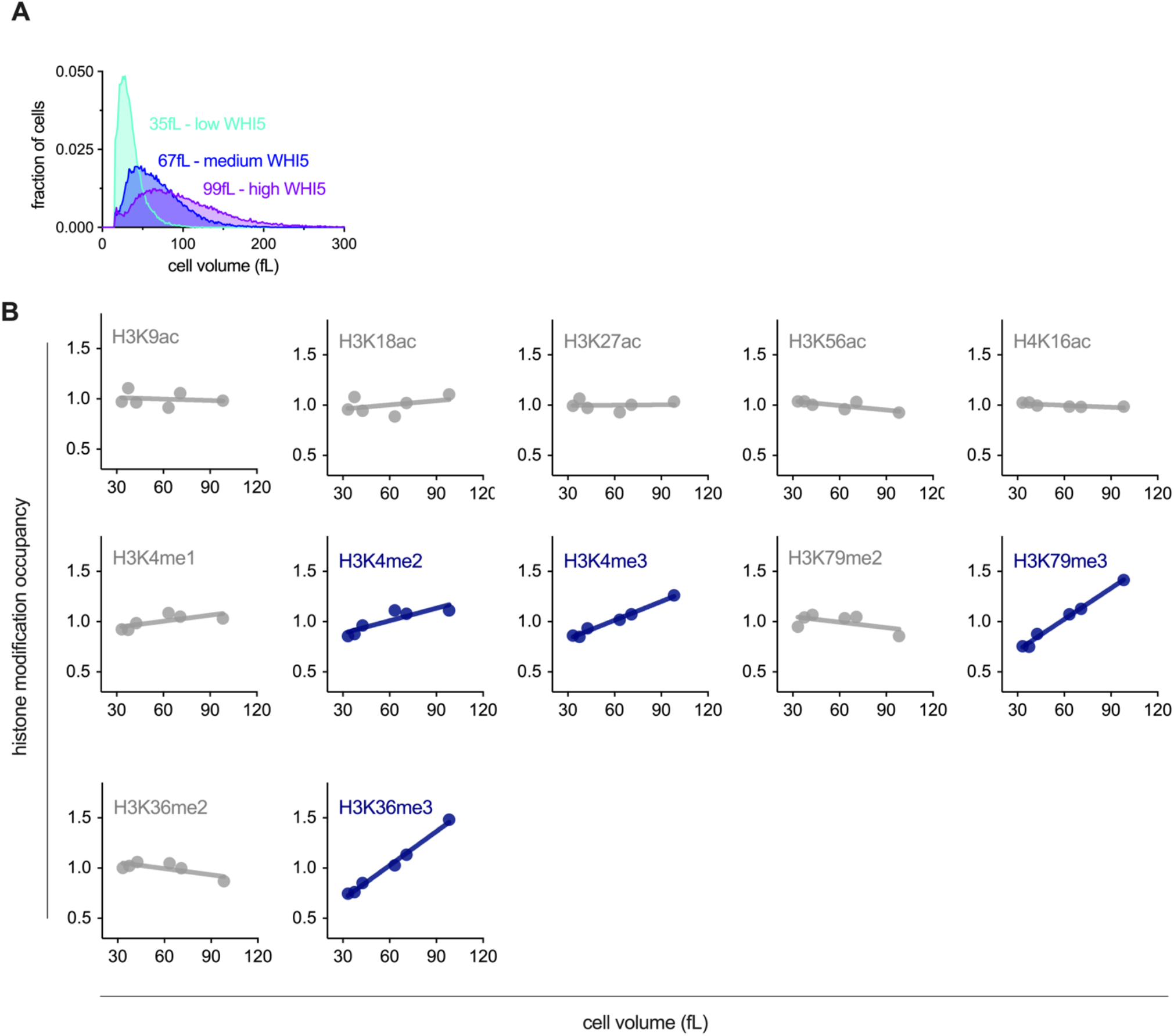
Analysis of chromatin as a function of cell size supporting data. Related to Fig. 5 **(A)** Cell size distributions determined by Coulter counter for the cultures used in the dSMF experiment in Fig. 5A-B to measure chromatin accessibility in cells of different sizes. **(B)** Global histone modification occupancy of the data shown in Fig. 5E. H3K56ac and H3k36me3 data are also shown in Fig. 5E. Line shows the linear fit used to calculate the slope values shown in Fig. 5E.

**Figure S8.**
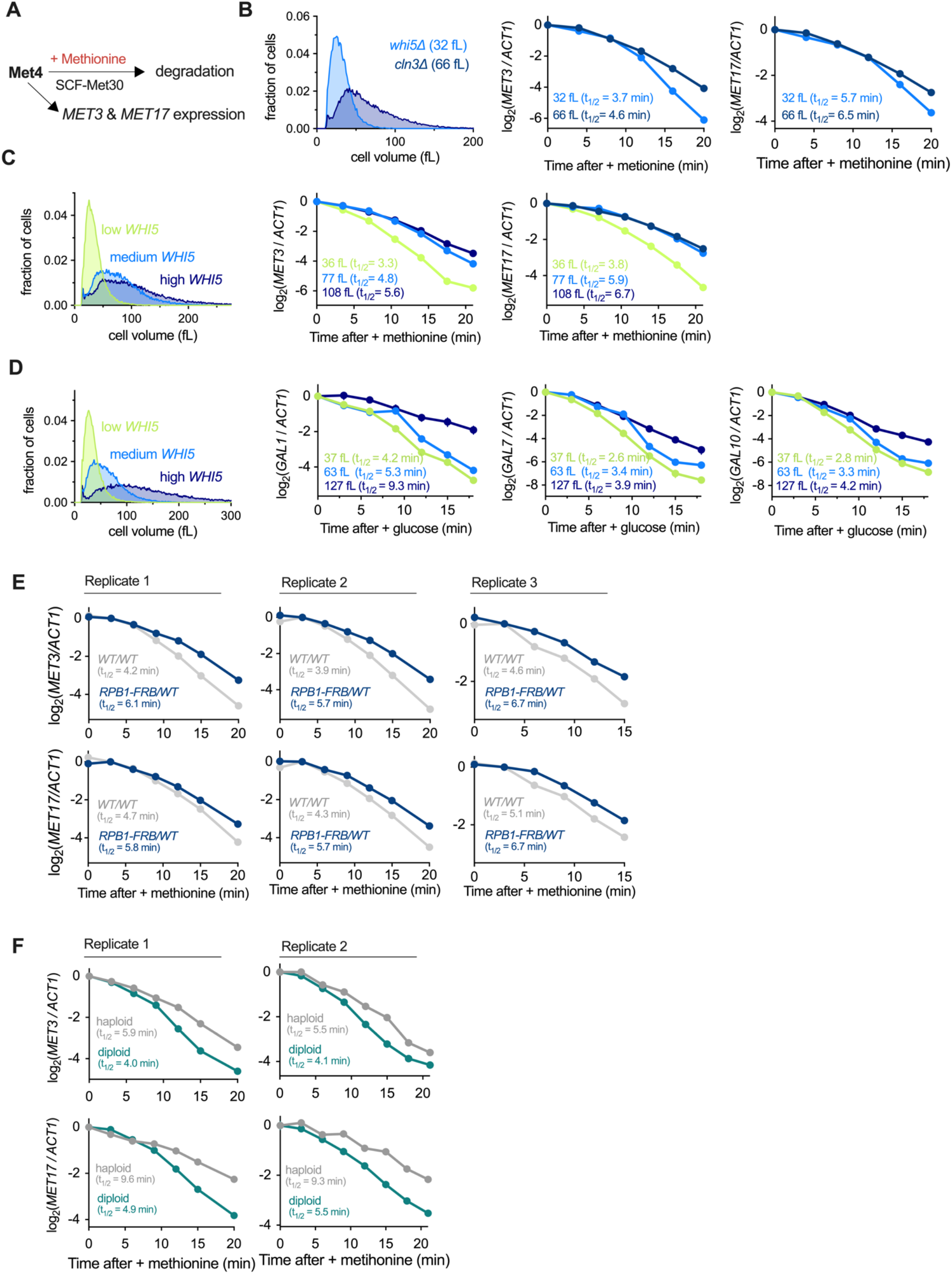
mRNA decay compensation supporting data. Related to Fig. 6 **(A)** Schematic showing methionine responsive repression of *MET* genes including *MET3* and *MET17* (Rouillon et al., 2000). **(B-C)** Example cell size distributions (left panel), *MET3* and *MET17* mRNA levels (centre and right panels) after transcriptional inactivation by methionine addition to cells of different sizes. mRNA levels are determined by rt-qPCR normalised to *ACT1*. Decay rate estimates from the one phase exponential decay fits to these data are shown in Fig. 6C. All experiments were performed in biological replicate. **(D)** Example cell size distributions (left panel), *GAL1, GAL7* and *GAL10* mRNA levels (centre and right panels) after transcriptional inactivation by galactose wash and glucose addition in cultures of different sizes. mRNA levels are determined by rt-qPCR normalised to *ACT1*. Decay rate estimates from the one phase exponential decay fits to these data are shown in Fig. 6C. **(E)** *MET3* (top) and *MET17* (bottom) mRNA levels relative to *ACT1* mRNA after transcriptional inactivation by methionine addition in *WT/WT* diploids or *RPB1-FRB/WT* heterozygous diploids. Methionine was added 40 minutes after rapamycin treatment. Rapamycin treatment is to deplete 50% RNAPII from the nucleus. The bottom middle panel is also shown in Fig. 6D. **(F)** *MET3* (top) and *MET17* (bottom) mRNA levels relative to *ACT1* mRNA after transcriptional inactivation by methionine addition in haploids or diploids arrested in G1 to ∼100fL by G1 cyclin shutoff. The bottom middle panel is also shown in Fig. 6E.

**Table S1.** List of *S. cerevisiae* strains and their genotypes used in this study

## Author contributions

MPS designed, performed and analysed all experiments except for the single molecule tracking experiments, which were performed and analysed by HZ and RRL, and the proteomics data acquisition performed by JG and AJ. Sequencing data analysis was performed by GM and MPS. JMS wrote the dynamic equilibrium model. MPS and JMS wrote the manuscript.

## Acknowledgments

We would like to thank Christine Jacobs-Wagner, Jim Ferrell, Kurt Schmoller, Mart Loog and members of the Skotheim lab for useful discussion and feedback on the manuscript. This work was supported by the NIH (GM092925 and GM115479), the HHMI-Simons (JMS, Faculty Scholars Program). MPS was supported by a Simons Foundation Fellowship of the Life Sciences Research Foundation and an EMBO Long-Term Postdoctoral Fellowship.

## Materials and methods

### Yeast genetics

All *S. cerevisiae* budding strains used in this study are in the *W303* background. Standard procedures were used for *S. cerevisiae* and *S. pombe* strain construction. Full genotypes of all strains used in this study are listed in **Table S1**. Diploid strains were constructed by crossing the respective parent haploid strains listed in **Table S1** and then streaking out on SCD -LEU -TRP selection plates.

Yeast strains MS627-635, used for the conditional overexpression of individual mNeonGreen tagged RNA polymerase II subunits (**Fig. 2E & S5**) were constructed as follows. pMS153, which encodes the WTC846 TET transcription factor system described in Azizoğlu et al., was linearized and integrated into the *HIS3* locus (Azizoğlu et al., 2020). Then each RNA polymerase II subunit was separately sub-cloned with a C-terminal mNeonGreen tag into the *URA3* SIV backbone (Wosika et al., 2016) downstream of the WTC846 *TETpr* (Azizoğlu et al., 2020) and upstream of the *AHD1* terminator. Each plasmid (pMS200-pMS208) was then linearized and integrated into the *URA3* locus. 3 out the 12 RNA polymerase II subunits where not tested (*RPB2*, *RPB5* and *RPB12*) because C-terminal tagging of the endogenous gene locus was either inviable or had a significant growth defect.

Yeast strain MS699, used for the simultaneous conditional overexpression of all RNA polymerase II subunits (**Fig. 2F-G**), was constructed as follows. First a plasmid (pMS152) which encodes the WTC846 TET transcription factor system described in Azizoğlu et al., was integrated into the HO locus (Azizoğlu et al., 2020). Four plasmids (pMS226, pMS230, pMS231 and pMS232) were then constructed, each based on a different single integration vector (SIV) backbone (Wosika et al., 2016) with three different RNA polymerase II subunits cloned in tandem into the into the multi-cloning site with an upstream WTC846 *TETpr* (Azizoğlu et al., 2020) with a downstream *ADH1* terminator. The *RPB3* construct (in pMS226) was also fused via a linker to the mNeonGreen fluorescent protein. Each plasmid was linearized and sequentially integrated into the *URA3*, *HIS3*, *TRP1* and *LEU2* loci, respectively, of MS46 and then *WHI5* was deleted. A control strain (MS656) was constructed in the same manner as MS669 but using four integration vectors each encoding mNeonGreen expressed from the WTC846 *TETpr* with an *ADH1* terminator downstream (pMS198, pMS233, pMS234 and pMS235).

### Yeast culturing conditions

Cells were grown in synthetic complete media (SC) with either 2% glucose, 2% glycerol + 1% ethanol or 2% raffinose as a carbon source. SC + 2% glycerol + 1% ethanol was used for all experiments examining cell size mutants (**Fig. 1E-F, 2B, S2-3, 3B-E & 6B-C)**, as this increases the dynamic range in size between mutants, apart from the *GAL* induction and shutoff experiments where cells where grown SC + 2% raffinose (**Fig. 4C** & **S8D;** see *qPCR mRNA decay experiments* section below). SC + 2% glycerol + 1% ethanol was also used for the elutriation experiments (**Fig. 3A-B, 4D-E & S6,** See *centrifugal elutriation* methods section below) and G1 arrest experiments (**Fig. 6E & S7**). All other experiments were performed in SC + 2% glucose. Unless stated otherwise, cells were grown at 30°C, always cultured below OD_600_ = 0.45, and collected at an OD_600_ between 0.2 and 0.4.

Low, medium and high expression of *WHI5* from MS63 (**Fig. 1D**), which carries an integrated copy of a *LexOPr-WHI5* construct, was induced by growing cells for >36 hours in 0nM, 10nM, or 30nM beta-estradiol, respectively, at which point cell size is at a steady state. Rapamycin was used at a final concentration of 1 µg/ml. Cells were collected 100 minutes (all samples in **Fig. S4B** and all samples in **Fig. 2D** except *Rpb1-FRB / WT*) or 40 minutes (*Rpb1-FRB / WT* samples in **Fig. 2D** and all samples in **Fig. S4C-H**) after rapamycin treatment. The shorter, 40 minute, rapamycin treatment was used due to compensation in Rpb1 protein amounts following 50% nuclear depletion that starts around 100 minutes following rapamycin treatment. Anhydrotetracycline was used at a final concentration of 50 ng/ml. Drug treatment duration is as specified in associated figure legends.

For the SILAC G1 arrest sample in **Fig. 2B & S3E,** MS54 was grown in the presence of 20nM beta-estradiol, which induces expression of *CLN1* and is essential for proliferation in this strain background lacking the other G1 cyclins (*cln2Δ cln3Δ*). To initiate the G1 arrest, cells were washed on a filter membrane (3x) in pre-warmed media lacking beta-estradiol and then returned to media lacking beta-estradiol.

For all SILAC experiments (**Fig. 2B & S3C-E**) SILAC compatible strains (*i.e.*, the *lys1Δ arg4Δ CAN1+* background) were grown for more than 10 generations in SILAC media supplemented with light (L-Arginine (unlabelled) & L-Lysine (unlabelled)), medium (L-arginine:HCL [U13C6] and L-lysine:2HCL [4,4,5,5-D4], or heavy (L-arginine:HCL (13C6, 15N4) and L-lysine:2HCL (13C6, 15N2)) amino acid isotopes (Cambridge Isotope Laboratories Inc.).

### Cell size measurements

Cell volume was measured using a Beckman Coulter Z2 counter. Cells were placed on ice, sonicated and then diluted in 10-20 ml of Isoton II diluent (Beckman Coulter #8546719) before measurement.

### DNA-content analysis

DNA content was determined by flow cytometry and was used to estimate genome copy number per cell for cell size mutants. 0.4 ml culture was added to 1 ml 100% 4°C ethanol and stored at 4°C. Cells were pelleted (13 krpm, 2 minutes), washed, and resuspended in 50 mM Sodium Citrate (pH = 7.2). They were then incubated with 0.2 mg/ml RNAse A (overnight, 37 °C) and then treated with 0.4 mg/ml proteinase K (1-2 hour, 50°C) before addition of 25 μM Sytox Green (ThermoFisher Scientific). Cells were then sonicated and DNA-content was analysed for >10,000 events on either a BD Biosciences FACScan Analyzer or an Attune NxT flow cytometer. Genome copy number per cell was then calculated after gating for single cells (FlowJo) and was used to convert occupancy per genome determined by spike-in normalized ChIP-seq to occupancy per cell (see *spike-in normalized ChIP-seq* methods section for details).

### Live cell flow cytometry measurements

Protein levels of C-terminally mNeonGreen-tagged proteins (**Fig. 2A**, **2E & S5**) were quantified using an Attune NxT flow cytometer. Strains were gently sonicated and placed on ice before acquisition of data from >10,000 cells per sample. Single cells were gated based on FSC and SCC (FlowJo). mNeonGreen intensity was measured in the BL1-A channel and cell size was approximated using FSC-A. The size-dependent autofluorescence background in the BL1-A channel was determined by measuring an untagged background control strain and fitting a robust linear regression to BL1-A vs FSC-A. This was then used to subtract the interpolated BL1-A background for each cell based on its FSC-A.

### Centrifugal elutriation G1 arrest experiment

For the G1 arrest elutriation experiment (**Fig. 3A-B, 4D-E & S6**), a G1 cyclin shut-off strain (*cln2Δ cln3Δ BEpr-CLN1*: MS64 (haploid) or MS67 (diploid)) was grown in SC + 2% glycerol + 1% ethanol with 15 nM beta-estradiol. SC + 2% glycerol + 1% ethanol media was used as this allow for the collection of the smallest G1 cells and does not require media carbon source removal or cooling to 4 °C during elutriation. The beta-estradiol is to induce expression of *CLN1* from a synthetic beta-estradiol responsive promoter. Small G1 cells arrested in the cell cycle where then collected as follows. 2 litres of culture were grown at 30°C to OD_600_ ∼0.7 and then washed twice on a filter membrane in fresh pre-warmed media lacking beta-estradiol and re-inoculated into 4 litres of media lacking beta-estradiol at 30 °C and grown for 2 hours. Cells were then collected on a filter membrane and resuspended in 100 ml fresh room temperature media. Cells were then sonicated (3 x 20 seconds, 4 minutes between sonication cycles) and loaded into a JE 5.0 elutriation rotor fitted for a two-chamber run (Beckman Coulter) in a J6-MI Centrifuge (2.4 krpm, 23 °C). The elutriation chambers were pre-equilibrated and run with room temperature SC +2% glycerol +1% ethanol media. The pump speed was gradually increased until small G1 cells with minimal debris were collected. The smallest G1 fractions were then combined and concentrated on a filter membrane, resuspended at 30°C in SC + 2% glycerol +1% ethanol media, and split into 7 cultures. Each culture was arrested in G1 for different periods of time before fixation and collection for ChIP-seq. At the point of splitting into 7 cultures, the OD_600_ of each culture was adjusted, according to its arrest time, so that each culture was at OD_600_ ∼0.3 at the point of fixation and collection. For both the haploid and diploid experiment two independent biological replicates was performed. A control experiment was also conducted where a *WT* strain (*i.e.*, not the G1 cyclin shut-off strain) was subjected to the same procedure, but after elutriation all cells were recovered from the elutriation rotor resulting in an asynchronous population that does not increase in cell size. Samples were then collected at the same timepoints used for the G1 arrest and therefore controls for all possible artefactual effects associated with handling of the cultures during elutriation.

### G1 arrest experiment

For the G1 arrest experiment without elutriation (**Fig. 6E & S8F**), a G1 cyclin shut-off strain (*cln2Δ cln3Δ BEpr-CLN1*: MS64 (haploid) or MS67 (diploid)) was grown in SC + 2% glycerol + 1% ethanol with 15 nM beta-estradiol. Cells were grown to OD_600_ ∼0.3 and washed 3x on a filter membrane in pre-warmed media lacking beta-estradiol to inhibit G1 cyclin expression and initiate the G1 arrest. Cells where then collected when the average cell volume reached approximately 100 fL.

### Microscopy

Cells were imaged on a wide-field epifluorescence Zeiss Observer Z1 microscope (63X/1.4NA oil immersion objective and a Colibri LED module). A single *z*-stack was taken, and cell boundaries were identified using phase contrast images. mCitrine fluorophores were imaged in the yellow channel (505 nm LED module). For the images in **Fig. S4A,** the exposure and contrast were manually adjusted in imageJ individually for each image to allow comparison of the relative subcellular localisation of the signal and is therefore not appropriate for comparing relative total protein intensities.

### Spike-in normalized ChIP-seq

The spike-in normalized ChIP-seq protocol was adapted from Hu *et al*. (Hu et al., 2015). *S. cerevisiae* cultures were combined with spike-in cultures (*C. glabrata* or *S. pombe*), mixed, and then immediately fixed (< 5 seconds after mixing) by the addition of formaldehyde to a final concentration of 1% (15 minutes). Cells were then quenched with 0.125 M glycine (5 minutes), washed twice in cold PBS, pelleted, snap-frozen, and stored at −80 °C.

Sample and spike-in were mixed in at a ratio between 1:2 and 1:5 by OD_600_. For most experiments *C. glabrata* (grown in the same media conditions as *S. cerevisiae*) was used as the spike-in. For experiments where anti-FLAG ChIP was performed, S. *pombe* with a Rpb1-3xFLAG (grown in EMM4S) was used as the spike-in. Within each batch of samples, the same spike-in culture was used for all samples and all samples were collected at effectively the same time (a 20-30 second interval between each sample and no more than 8 samples were collected in each batch). The only exception to the above is for the elutriation G1 arrest experiments where different *S. cerevisiae* cultures could not be mixed with the spike-in at the same time (because different samples required different arrest durations). For this experiment *S. cerevisiae* and *C. glabrata* were therefore separately fixed, quenched, washed, and then mixed in PBS before being pelleted and snap frozen.

For the experiments in **Fig. 2-4**, a background sample was included within each batch of experiments to determine the *S. cerevisiae* background-to-spike-in ratio. For anti-Rpb1 ChIP, the background was determined using a sample where TBP was conditionally depleted from the nucleus (MS205 or MS289; *TBP-FRB* treated with rapamycin (60-65 minutes rapamycin for SC + 2% glucose, 90 minutes rapamycin for SC + 2% glycerol + 1% ethanol) to block Rpb1 loading on the genome. For anti-FLAG ChIP, the background was determined using an untagged sample with no FLAG epitope (MS207). Background samples were otherwise collected and processed as other samples. See *Spike-in normalized ChIP-seq analysis* section for how this is used to calculate the global occupancy value for each ChIP.

Pellets were thawed and lysed in 300 µl FA lysis buffer (50 mM HEPES–KOH pH 8.0, 150 mM NaCl, 1 mM EDTA, 1% Triton X-100, 0.1% sodium deoxycholate, 1 mM PMSF, Roche protease inhibitor) with ∼1 ml ceramic beads on a Fastprep-24 (MP Biomedicals). The entire lysate was then collected and adjusted to 1 ml before sonication with a 1/8” microtip on a Q500 sonicator (Qsonica) for 16 minutes (cycles of 10 seconds on and 20 seconds off). The sample tube was held suspended in a −20°C 80% ethanol bath to prevent sample heating during sonication. Cell debris was then pelleted, and the supernatant retained for ChIP or input. For each ChIP reaction, 20 µl Protein G Dynabeads (Invitrogen) were blocked (PBS + 0.5% BSA, incubate 40 minutes at room temperature), pre-bound with 5 µl of antibody in PBS (incubate 40 minutes at room temperature), and washed 2x with PBS before being incubated with 0.5 ml supernatant (4°C overnight). For RNA polymerase II ChIPs, the supernatant was adjusted so 0.5 ml corresponds to 50-100 ml of cells at OD_600_ ∼ 0.35. For histone and histone modification ChIPs, the supernatant was adjusted so 0.5 ml corresponds to 10-25 ml of cells at OD∼0.35. See below for a list of antibodies used. After overnight incubation, Dynabeads were washed (5 minutes per wash) 2x in FA lysis buffer and 3x in high-salt FA lysis buffer (50 mM HepesKOH pH 8.0, 500 mM NaCl, 1 mM EDTA, 1% Triton X-100, 0.1% sodium deoxycholate, 1 mM PMSF). ChIP DNA was then eluted in ChIP elution buffer (50 mM TrisHCl pH 7.5, 10 mM EDTA, 1% SDS) at 65°C for 20 minutes. At the same time, 15 µl of input was mixed directly with 115 µl of ChIP elution buffer. Eluted ChIP DNA and input DNA were then incubated to reverse crosslinks (65°C, 5hr), before treatment with RNAse A (37 °C, 1 hour) and then Proteinase K (65°C, 2 hours). DNA was then purified using the ChIP DNA Clean & Concentrator kit (Zymo Research). Indexed sequencing libraries were generated using the NEBNext Ultra II DNA Library Prep kit (NEB Cat # E7645), pooled, and then sequenced by paired-end (2×150bp) Illumina sequencing.

The following antibodies were used for ChIP against the indicated epitopes: anti-Rpb1 clone 8wG16 (mouse, monoclonal, Millipore # 05-952-I), anti-Rpb1-S2-P clone 3E10 (rat, monoclonal, Millipore, #04-1571-1), anti-Rpb1-S5-P clone 3E8 (rat, monoclonal, Millipore #04-1572-I), anti-FLAG clone M2 (mouse, monoclonal, Sigma-Aldrich #F3165), anti-Histone 3 (rabbit, polyclonal, Abcam #ab1791), anti-Histone 4 (rabbit, polyclonal, Abcam #ab10158), anti-H3K18ac (rabbit, polyclonal, Abcam #ab1191), anti-H3K27ac (rabbit, polyclonal, Millipore #07-360), anti-H4K9ac (rabbit, polyclonal, Abcam #ab4441), anti-H4K16ac (rabbit, polyclonal, Millipore #07-329), anti-H3K56ac (rabbit, polyclonal, Millipore #07-677), anti-H3K4me1 (rabbit, polyclonal, Abcam #ab8895), anti-H3K4me2 (rabbit, polyclonal, Abcam #ab7766), anti-H3K4me3 (mouse, monoclonal, Abcam #ab1012), anti-H3K79me2 (rabbit, polyclonal, Abcam #ab3594), anti-H3K79me3 (rabbit, polyclonal, Abcam #ab2621), anti-H3K36me2 (rabbit, polyclonal, Abcam #ab9049), anti-H3K36me3 (rabbit, polyclonal, Abcam #ab9050).

### Spike-in normalized ChIP-seq analysis

Unless otherwise specified, all analyses described were carried out using custom-written Python scripts (https://github.com/georgimarinov/GeorgiScripts).

Combined *S. cerevisiae* and spike-in (*C. glabrata* or *S. pombe*) genome FASTA files were created using the sacCer3 genome assembly for *S. cerevisiae* and the assemblies for *C. glabrata* (obtained from the Candida Genome Database (CGD) (Skrzypek et al., 2017)) or *S. pombe (*ASM294v2). Combined gene annotations were created using *S. cerevisiae* gene models updated using transcript-end mapping data as previously described (Shipony et al., 2020) and either the *C. glabrata* gene models available from CGD or the *S. pombe* gene models available from PomBase (Lock et al., 2019).

Demultipexed FASTQ files were then mapped to the relevant combined genome indexes as 2×36mers using Bowtie (v.1.0.1) (Langmead et al., 2009) with the following settings: -v 2 -k 2 -m 1 --best --strata. For subsequent analyses, reads that map to the *S. cerevisiae* or spike-in mitochondrial genomes were ignored and reads that map to both the *S. cerevisiae* and spike-in genomes were ignored.

To determine the global occupancy ratio, the sample-to-spike-in ChIP ratio of reads was calculated and then divided by the sample-to-spike-in input ratio for the same sample. The global occupancy ratio is therefore expressed per genome. For the data in **Fig. 1E** the occupancy per genome was converted to occupancy per cell by multiplying by the average genomes per cell determined by flow cytometry (see *DNA content analysis* section above). The Histone modification occupancy (**Fig. 4C-E & S7B-C**) was calculated as above and then normalized to the anti-H3 or anti-H4 global occupancy ratio for the same sample. For the anti-Rpb1 and anti-FLAG ChIP experiments in **Fig. 2, 3, 4, S4 & S6** the *S. cerevisiae* background-to-spike-in ratio was determined (see *Spike-in normalized ChIP-seq* section above for details on what samples were used to determine the background ratio). For these data, the background ratio was then subtracted from the global occupancy ratio. For the G1 arrest ChIP data in **Fig. 3 & 6** the global occupancy ratios were then normalised to occupancy ratios of asynchronous control samples that were collected at the same timepoints but where not arrested in G1.

RPM (Reads Per Million) normalized read coverage genome browser tracks showing the 50bp mid-point of each mapped fragment (*i.e.*, midpoint ± 25bp) were generated for the *S. cerevisiae* genome using custom-written python scripts. Subsequent analysis uses these 50bp mid-point read coverage tracks to calculate per gene occupancies and metagene profiles. Per gene occupancy was calculated as follows. RPKM values were calculated for the gene body (TSS-to-TTS) for all verified ORFs. ChIP RPKM values were then multiplied by the global ChIP occupancy factor described above to give the per gene occupancy value. Metagene profiles showing the average occupancy across gene bodies were generated as follows. Average RPKM values were calculated for the 500bp upstream of the TSS, 500bp downstream of the TTS and gene bodies (TSS-to-TTS; rescaled to be 1kb) and then multiplied by the global occupancy factor described above and were smoothed using a 50bp averaging window. The top 10% of genes (verified ORFs only) was determined based on anti-Rpb1 RPKM ChIP and was used where indicated. For the histone modification metagene heatmaps (**Fig. S7B**), values were re-scaled to the median value of all samples displayed on a given heatmap to aid comparison between modifications.

### Dual enzyme single-molecule foot printing (dSMF)

The protocol for dSMF to measure chromatin accessibility (**Fig. 5A-B**) was based on Krebs *et al*., (2017) (Krebs et al., 2017) and adapted for yeast as follows. 1×10^8^ *S. cerevisiae* cells were pelleted and resuspended in 1 ml digestion buffer (1.4 M sorbitol, 40 mM Hepes-KOH[pH7.5], 0.5 mM MgCl_2_, 10 mM DTT) and mixed with 1×10^7^ *C. glabrata* cells (spike-in) in 100 µl digestion buffer. Cells were pelleted and resuspended in 0.5 ml digestion buffer + 0.5 mg/ml 100T Zymolase (MP biomedicals), and then incubated on a shaker (30°C, 10 minutes, 500 rpm). Cells where then pelleted (5k rpm, 2 minutes, 4°C), resuspended in 0.5 ml wash buffer (1.4 M sorbitol, 40 mM Hepes-KOH[pH7.5], 0.5 mM MgCl_2_), pelleted again (5k rpm, 2 minutes, 4°C), resuspended in 0.3 ml ice-cold lysis buffer (10 mM Tris-HCl[pH7.5], 10 mM NaCl, 3mM MgCl_2_, 0.1mM EDTA[pH7.5], 0.5% NP-40), and incubated on ice (10 minutes). Nuclei were then pelleted (5k rpm, 4 minutes, 4 °C), resuspended in 0.3 ml nuclei wash buffer (10 mM Tris-HCl[pH7.5], 10 mM NaCl, 3mM MgCl_2_, 0.1mM EDTA[pH7.5]), pelleted again, and resuspended in 50 µl 1x reaction buffer (50 mM Tris-HCl[pH8.5], 50 mM NaCl, 10mM DTT). Nuclei were then treated with M.CviPI (NEB) and M.SssI (NEB) methyltrasferases as follows. 50 µl cells in M.CviPI reaction buffer were mixed with 47 µl M.CviPI reaction mix (50 mM Tris-HCl[pH8.5], 50 mM NaCl, 10 mM DTT, 0.7 M sucrose, 1.3 mM SAM, 200 U M.CviPI) and incubated (8 minutes, 30°C) before mixing with 3 µl M.CviPI boost mix (100 U M.CviPI, 42.5 µM SAM), and returning to incubation (7 minutes, 30 °C). 10 µl M.SssI reaction mix (50 mM Tris-HCl[pH8.5], 50 mM NaCl, 10 mM DTT, 0.11 M MgCl_2_, 12.8 µM SAM, 60 U M.SssI) was then added to the 100 µl M.CviPI reaction, mixed and incubated (8 minutes, 30°C). The reaction was then stopped with the addition of 190 µl lysis buffer (from MasterPure Yeast DNA Purification Kit (Lucigen)), and DNA was extracted with a MasterPure Yeast DNA Purification Kit (Lucigen). Unmethylated cytosines were then deaminated to uracil and indexed libraries were prepared using the NEBNext Enzymatic Methyl-seq Kit (NEB #E7120) before being pooled and then sequenced by Illumina paired-end (2×150bp) sequencing.

### Dual enzyme single-molecule foot printing (dSMF) data analysis

Combined *S. cerevisiae* and *C. glabrata* genome and transcriptome files were created as described above (see *Spike-in normalized ChIP-seq analysis*). FASTQ files were trimmed of adapter using cutadapt (version 0.16) and Trim Galore (version 0.4.4) with the following settings: --clip_R1 9 --clip_R2 9 --three_prime_clip_r1 6 --three_prime_clip_r2 6 --paired. Trimmed reads were then mapped to the combined genome indexes using the bwameth package (https://github.com/brentp/bwa-meth). Duplicate reads were removed using the MarkDuplicates programs (picard-tools-1.99). Methylation calls were extracted using the MethylDackel (https://github.com/dpryan79/MethylDackel) with the following settings: --CHG --CHH. The MethylDackel output was used to create genome browser tracks showing methylation levels and to make metagene plots which were then normalized to the global methylation fraction determined for the spike-in *C. glabrata* genome.

### qPCR mRNA decay experiments

To quantify *MET3* and *MET17* turnover, cells were grown in media lacking methionine and 1 mM methionine was then added to repress MET genes (Rouillon et al., 2000). To quantify *GAL1*, *GAL7* and *GAL10* decay rates, cells were grown in SC media + 2% raffinose before addition of 2% galactose for 75 minute and then washed (1x) and resuspended in SC + 4% glucose (Haimovich et al., 2013). 1 ml of culture was pelleted (13 krpm, 30 seconds, 4 °C) at the indicated time points after either methionine addition (for *MET3* and *MET17*) or after the start of the wash in SC + 4% glucose (for *GAL1*, *GAL7* and *GAL10*). The pellet was immediately snap frozen in liquid nitrogen at t + 1 minute and stored at -80°C. Cells were subsequently thawed in 300 µl TRI Reagent (Zymo Research) and lysed by bead beating using a Fastprep 24 (4 °C, settings: 5.0 m/s, 1 x 30 seconds). Cell debris was pelleted (13 krpm, 2 minutes) and the supernatant recovered. RNA was then purified using the direct-zol RNA microprep kit (Zymo Research #R2061). cDNA was then synthesized using 800ng of RNA with iScript Reverse Transcription Supermix for RT-qPCR (BioRad #1708841) and used for qPCR for *MET3*, *MET17* and *ACT1* using iTaq Universal SYBR Green Supermix (BioRad #1725121). 2-3 biological replicates were performed per experiment and 2-4 technical replicates were performed per biological replicate. Decay rates (**Fig. 6C-E**) were calculated from a one-phase exponential decay fit to the data for timepoints up to 20 minutes after treatment (Prism 6; weighting = 1/Y). 0-minute timepoints (collected before methionine or glucose addition) were excluded from the calculation assuming a time-delay before inhibition starts, which also resulted in an improved fit.

### Single molecule imaging: microscopy

1 ml culture (SC + 2 % glucose) was pelleted (4k rpm, 1 minute) and resuspended in 0.5 ml fresh media. JF-PA549 (Janelia Farms photoactivatable 549) dye was added at a final concentration of 75 nM, except for MS162 (HTB1-HALO), where a concentration of 10 nM was used to compensate for the higher protein copy number. Cultures were incubated with dye (30°C, mixing at 550 rpm) for 40 minutes. Cells were then washed (3x) in fresh media to remove unbound dye and resuspended in 20 µl media, 4 µl of which was placed on an agarose pad. The agarose pad was made by mixing 0.5 ml 2% agarose Optiprep mixture (20 mg agarose in 1 ml Optiprep (Sigma), heated to 90 °C) with 0.5 ml 2x media. Approximately 110 µl of this mixture was placed within a Gene Frame (Thermo Scientific), with excess being removed with a KimWipe. Prior to imaging, we waited ∼15 minutes to let any remaining unbound dye be released from cells. Coverslips were cleaned using 2% VersaClean detergent solution overnight. The coverslips were then washed with MilliQ water 3 times, sonicated in acetone for 30 minutes, washed with MilliQ water 3 times, washed in methanol (flame excess from coverslips), and then placed in a Plasma Etch plasma oven for 10 minutes.

Imaging was done at 23°C using a Leica DMi8 inverted microscope with a Roper Scientific iLasV2 (capable of ring total internal reflection fluorescence (TIRF)) and an Andor iXon Ultra 897 EMCCD camera. An Andor ILE combiner was used and the maximum power from the optical fiber was 100 mW for the 405 nm wavelength, and 150 mW for the 488 nm and 561 nm wavelengths. The iLasV2 was configured for HILO (ring highly inclined and laminated optical sheet), selective illumination and single-molecule sensitivity. Metamorph software was used to control acquisition. A Leica HCX PL APO 100x/1.47 oil immersion objective was used with 100 nm pixel size. Z-stacks were determined using a PIano piezo Z controller. Single-particle photoactivated localization microscopy (sptPALM) experiments were performed by using continuous activation of molecules with low power (0.1% – 10% in Metamorph software) 405nm light to photo activate ∼1 molecule/cell, with simultaneous fast-exposure (10 ms) illumination with 561 nm light (70% in Metamorph software) to image molecules. A bright field and a 561 nm z-stack of 10 µm (0.5 µm step size) were taken and used to identify the unbudded G1 phase cells, quantify nuclear area using the Pus1-GFP nuclear marker. Nuclear area can then be used to estimate cell volume.

### Single molecule imaging: analysis

Aside from tracking, all analysis was performed using custom matlab code. Tracking was performed using Trackmate (Tinevez et al., 2017). First, molecules were localized in each frame using a Laplacian of Gaussian (LoG) method with an estimated diameter of 5 pixels. An intensity threshold was chosen that was low enough to still detect molecules that were moving out of the focal plane and were diffusing quickly. After localization, tracks were formed using the Linear Assignment Problem algorithm by linking molecules in consecutive frames. The linking distance was set to 5 pixels. A gap frame of 3 was used to allow for missed localization. The gap-linking distance was set to 5 pixels more than the linking distance. Linking also had a cost of 0.3 for the ‘‘Quality’’ parameter to ensure that correct molecules were linked. Tracks with fewer than five localizations were discarded.

Only unbudded G1 cells, determined from brightfield images, were retained for subsequent analysis. The cell outlines were segmented by a custom-made matlab code and then manually curated using the “Freehand” function in matlab. Cell volumes were estimated from the cell masks assuming ellipsoid geometry. The volumes of cells were estimated by adding up the cross-section volume at each orthogonal pixel layer of the major axis of cells. Maximum intensity projections of z-stacks in 488nm were then used for segmenting the nuclear regions using the Pus1-GFP nuclear marker and tracks outside the nucleus were discarded. The radius of gyrations (RoG) was then calculated for each track using the following equation:

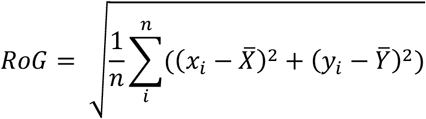

where n is the number of localisations in each track, and (x_i_, y_i_) are the coordinates of the track for each step i, and (*C̄*,*Ȳ*) are the mean of the x, y coordinates of the track, respectively. To classify molecules as bound or unbound, a one-dimensional Gaussian mixture model (GMM) was fitted to the log2-transformed RoG distribution. The initial values for fitting were inferred from the log-transformed RoG distribution of H2B1-HALO and mCitrine-HALO-NLS, which have the majority of molecules bound and unbound, respectively. To determine how many Gaussian groups exist in the RoG distribution, we fitted the distribution with different GMMs, where total gaussian groups varied from 1 to 4. The Bayesian information criterion (BIC) was calculated for each GMM fitting. The model with 2 gaussian groups was selected on the basis that it had the lowest BIC. The Gaussian group with the lowest average RoG value was classified as the group of bound molecules. The tracks inside the chromatin bound fraction in each nucleus was then determined by dividing the number of bound tracks by the number of total tracks. Nuclei with < 30 tracks were discarded from this analysis.

### Chromatin enrichment for Proteomics (ChEP)

For SILAC experiments, SILAC compatible strains were labelled with amino acid isotopes (see *Yeast culturing conditions* for details) before collection. The ChEP chromatin enrichment protocol was based on Kustatscher *et al*., (Kustatscher et al., 2014) and adapted for yeast as follows. Cells were fixed by addition of formaldehyde to a final concentration of 1%, shaken for 15 minutes, quenched with 0.125 M glycine (5 minutes), washed twice in cold PBS, pelleted, snap-frozen, and then stored at −80°C. Each pellet contained cells from 500 ml culture at OD_600_ ∼ 0.4. Pellets were thawed and lysed in 300 µl FA lysis buffer (50 mM HEPES–KOH pH 8.0, 150 mM NaCl, 1 mM EDTA, 1% Triton X-100, 0.1% sodium deoxycholate, 1 mM PMSF, 1x Roche protease inhibitor) with ∼1 ml ceramic beads on a Fastprep-24 (MP Biomedicals). The entire lysate was then collected and adjusted to 1.5 ml.

For the cytoplasmic fraction (CYTO), 50 µl lysate was adjusted to 0.5 ml, pelleted (4°C, 14k rpm, 10 minutes), and the supernatant was then taken as the cytoplasmic fraction. At this stage protein concentration was determined by Bradford before Laemmli sample buffer was added to a 1x final concentration and the sample was then boiled to reverse crosslinks (99 °C, 30 min). For whole cell extract (WCE), 50 µl lysate was adjusted to 0.5 ml and then sonicated with a 1/8” microtip on a Q500 sonicator (Qsonica) for 5 minutes (Amp = 25%, 10 seconds on, 20 seconds off) to solubilize chromatin. Laemmli sample buffer was added to a 1x final concentration, and the sample was then boiled to reverse crosslinks (99 °C, 30 min) before pelleting (4°C, 14k rpm, 10 minutes). The supernatant was then taken as the WCE. WCE protein concentration was assumed to be the same as was determined for the cytoplasmic fraction for the same sample.

For chromatin fractions the lysate was pelleted (4°C, 14k rpm, 30 minutes) and supernatant was removed and discarded. The pellet was retained and either processed with extraction method A (high purity, low yield) or extraction method B (high purity, low yield). For extraction method A the fuzzy translucent top layer (high purify chromatin) of 3 pellets was resuspended in 300 µL FA lysis buffer without disturbing the lower opaque portion of the pellets and combined to a single fresh tube and adjusted to 1.45 ml. For extraction method B, 1 entire pellet (top and bottom layer) was resuspended in FA lysis buffer to a final volume of 1.45 ml. Extractions A and B were then processed identically for all subsequent steps. Chromatin was first treated with 100 µg/ml RNaseA (37°C, 15 minutes) and then pelleted (4 °C, 14k rpm, 30 minutes). The supernatant was then discarded and the pellet was resuspended in 300 µL SDS buffer (4 % SDS, 10 mM EDTA, 25 mM TrisHCl pH 7.5, 1 mM PMSF, 1x Roche protease inhibitor) and incubated at room temp (10 minutes). 1 ml urea buffer (8M urea, 1mM EDTA, 10mM TrisHCl pH 7.5, 1 mM PMSF, 1x Roche protease inhibitor) was then added and the sample was gently mixed. The sample was then pelleted (18°C, 14k rpm, 30 minutes) and the supernatant was discarded. The pellet was resuspended in 300 µl SDS buffer mixed with 1 ml urea buffer before being again pelleted (18°C, 14k rpm, 30 minutes). The supernatant was discarded and the pellet was resuspended in 300 µl SDS buffer and then adjusted to 1.45 ml with SDS buffer and again pelleted (18°C, 14k rpm, 30 minutes). The supernatant was again discarded and 0.6 ml Storage buffer (10% glycerol, 1 mM EDTA, 10 mM TrisHCl pH 7.5, 25 mM NaCl, 1 mM PMSF, 1x Roche protease inhibitor) was added on top of the pellet. DNA was then resuspended and sheared by sonication (1/8” microtip on a Q500 sonicator (Qsonica) for 10 minutes (Amp = 25%, 10 seconds on, 20 seconds off)). The sonicated sample was then pelleted (4 °C, 14k rpm, 10 minutes) and the supernatant was retained as the chromatin fraction. Protein concentration was determined by Bradford before Laemmli sample buffer was added to a 1x final concentration and the sample was then boiled to reverse crosslinks (99°C, 30 min).

For the immunoblotting analysis presented in **Fig. S3B**, ∼ 5 µg of the chromatin fraction B was loaded per lane alongside ∼25 µg CYTO and WCE. For the proteomics analysis in **Fig. S3C-D**, chromatin and WCE with opposite SILAC labels were mixed in an approximately 1:2 protein amount ratio before processing ∼ 100 µg for LC-MS/MS (for details see *LC-MS/MS sample preparation and data acquisition* below). For the proteomics analysis in **Fig. 2B**, heavy, medium and light labelled cultures were mixed prior to fixation in an approximately 1:1:1 cell number ratio. Two biological replicates were performed for chromatin extraction B. ∼50 µg of each replicate was then analysed by LC-MS/MS (for details see *LC-MS/MS sample preparation and data acquisition*).

### Immunoblotting

Protein samples were resolved on a Bolt 4-12% Bis-Tris protein gel (invitrogen) and transferred to a nitrocellulose membrane with the iBlot 2 dry blotting system (Invitrogen). The following primary antibodies were used for western-blotting at 1/1,000 dilution: anti-V5 clone SV5-Pk1 (mouse, monoclonal, BioRad, #MCA1360), anti-Rpb3 clone 1Y26 (mouse, monoclonal, BioLegend #665004), anti-FRB (rabbit, polyclonal, Enzo #ALX-215-065-1), anti-Beta tubulin (rabbit, polyclonal, Abcam #ab15568), anti-Histone 3 (rabbit, polyclonal, Abcam #ab1791) and anti-Histone 4 (rabbit, polyclonal, Abcam #ab10158). Anti-GAPDH clone GAR1 (mouse, monoclonal, ThermoFisher #MA5-15738) was used for western-blotting at 1/2,500 dilution. Primary antibodies were detected using the following fluorescently labelled secondary antibodies at 1/10,000 dilution: IRDye 800CW goat anti-Mouse (Licor), Alexa Fluor 680 Donkey anti-Mouse (Invitrogen), Alexa Fluor 680 donkey anti-Rabbit (Invitrogen) and Alexa Fluor 790 Goat anti-Rabbit (Invitrogen). Membranes were then imaged using a LI-COR Odyssey CLx.

### LC-MS/MS sample preparation and data acquisition

Each protein sample was reduced with 5 mM dithiothreitol for 25 minutes at 56°C, alkylated with 10 mM iodoacetamide (30 minutes, room temperature, dark) and then quenched with 7.5 mM DTT. Samples were digested and cleaned using SP3 on-bead methodology (Hughes et al., 2019) with the variation that 50 mM HEPES (pH 8.5) was used in place of ammonium bicarbonate. Briefly, proteins were bound to the SP3 beads (10:1 beads:protein (w/w)) in 50% ethanol (v/v) and then washed three times in 80% ethanol, prior to resuspension in 50 mM HEPES (pH 8.5) with 1:40 (trypsin:protein (w/w)) overnight at 37°C. The peptides were then fractionated using the High pH Reversed-Phase Peptide Fractionation Kit (Pierce) and dried under vacuum centrifugation. Peptides were subsequently resuspended in 0.1% trifluoroacetic acid and analysed on an Orbitrap Fusion Lumos mass spectrometer (Thermo Fisher) coupled to an UltiMate 3000 HPLC system for online liquid chromatographic separation. Each run consisted of a 160-minute gradient elution from a 75 µm x 50 cm C18 column.

### Analysis of ChEP LC-MS/MS data

MaxQuant (version 1.5.0.13) was used for all LC-MS/MS data processing. The data were searched against a UniProt extracted *S. cerevisiae* proteome FASTA file. SILAC comparison between WCE and chromatin fractions (See **Fig. S3C-D** and methods section *Chromatin enrichment for Proteomics (ChEP)*) was used to determine a high confidence list of chromatin-enriched proteins (n=564). Chromatin-enriched proteins were defined as proteins enriched by chromatin extraction methods A and B compared to WCE in both biological replicates. These 564 proteins and the threshold used to define enrichment are shown in **Fig. S3D**. Normalization of SILAC ratios was not applied because there is no *a priori* assumption that the median of the distribution corresponds to no change. Instead, raw SILAC ratios are plotted and the thresholds for each replicate were defined so that thresholds intersect on the linear regression line between the two replicates (**Fig.S3D**) at a point that separates known chromatin associated factors from abundant cytoplasmic factors (**Fig.S3C**). The distribution of SILAC ratios for the experiment plotted in **Fig. 2B** was normalized so the average histone protein corresponds to a SILAC ratio = 1 and as such these data can be interpreted as protein bound per genome, given histone occupancy per genome is approximately constant as a function of cell size, as determined by histone Chip-Seq (**Fig. S6D**).

